# Cultivar-specific transposable element insertions are associated with both suppression and dominance of homoeolog expression in bread wheat

**DOI:** 10.64898/2026.11.24.768836

**Authors:** Liel Gribun, Khalil Kashkush

**Author notes:** Correspondence: Khalil Kashkush, Department of Life Sciences, Ben-Gurion University, Beer-Sheva 8410501, Israel.

## Abstract

Bread wheat is an allohexaploid in which the three homoeologous copies of most genes are expressed unequally, and transposable elements (TEs) occupying gene bodies have been proposed as one source of that imbalance. Previous tests of this idea examined a single genotype, so they could not ask whether differences in insertion between cultivars correspond to differences in expression. We sequenced young-leaf transcriptomes of six bread wheat cultivars, quantified them against IWGSC RefSeq v2.1, and combined them with transposable element annotations generated independently for each cultivar’s own genome. Between 29.47% and 29.64% of genes carried an element that lay entirely within the gene body, 85% of which were intronic. Across all 12,887 homoeologous groups analyzed, agreement between insertion and expression exceeded a permutation null only modestly. Restricting the test to the 1,692 groups whose insertion pattern differs between cultivars changed the picture: 115 of these showed expression bias tracking the insertion, against 27.3 expected, a 4.21-fold enrichment. The association was markedly asymmetric, reaching 6.83-fold for suppression of the inserted homoeolog but 2.53-fold for dominance, and it was carried by intronic rather than coding-sequence insertions and by six of eleven element superfamilies. The same trend was observed across all six tissues in an independently generated wheat pan-transcriptome, where quantifying the same reads against each cultivar’s own genome assembly confirmed the association independently of the reference used. We identified 74 candidate loci at which polymorphic insertions were associated with consistent transcriptional suppression across all six cultivars. These findings reveal polymorphic TE insertions as a source of evolutionary variation in homoeolog expression, linking genome structural diversity to transcriptional diversification in bread wheat.

**Significance statement:** TEs are found in almost a third of bread wheat genes, but whether they shape the unequal expression of homoeologous gene copies has been tested only within single genotypes. Comparing six cultivars, we show that expression bias tracks insertions specifically where they differ between cultivars, in both directions, but far more strongly towards suppression.

## Introduction

Bread wheat (*Triticum aestivum*, 2n= 6x= 42, AABBDD) is the product of two hybridization events that brought three related diploid genomes into a single nucleus. The A and B subgenomes were united less than half a million years ago in the tetraploid progenitor of wild emmer (*T. turgidum* ssp. *dicoccoides*), whose genome architecture and diversity have since been resolved in detail (Marcussen et al., 2014; Avni et al., 2017), and the D subgenome was contributed some 8,000 to 10,000 years ago by *Aegilops tauschii* (Luo et al., 2017). The resulting allohexaploid spread from the Fertile Crescent to become the most widely cultivated crop on Earth, and the genetic consequences of that history remain visible in the modern crop: tracing the ancestry of present-day bread wheats recovers distinct founder contributions to each subgenome (Pont et al., 2019). A chromosome-scale reference assembly of the cultivar Chinese Spring (International Wheat Genome Sequencing Consortium, 2018), subsequently refined with optical maps into RefSeq v2.1 (Zhu et al., 2021), and the assembly of fifteen further wheat genomes by the 10+ Wheat Genomes Project (Walkowiak et al., 2020) have together made it possible to study this 16-gigabase genome not as a single reference but as a population of genomes.

More than 85% of the wheat genome consists of transposable elements, dominated by long terminal repeat retrotransposons of the *Gypsy* and *Copia* superfamilies, with DNA transposons such as *CACTA* and a large complement of miniature inverted-repeat elements making up much of the remainder (Wicker et al., 2007, 2018). This compartment is not inert. Transcriptional activation of retrotransposons in newly formed wheat allopolyploids alters the expression of adjacent genes (Kashkush et al., 2003), and work across wheat species has since documented how elements continue to reshape the genome: miniature inverted-repeat elements are distributed non-randomly with respect to genes and show lineage-specific amplification (Keidar-Friedman et al., 2018, 2020), retroelements such as Au SINE generate population-specific insertional patterns and allelic variation (Domb et al., 2017), and element activity underlies large-scale structural rearrangement during wheat evolution (Bariah et al., 2020a, 2020b) and population-specific structural variation in contemporary germplasm (Bida and Kashkush, 2026). Critically for the present study, individual elements are frequently polymorphic between cultivars, to the point that their insertion sites serve as practical molecular markers for genotyping wheat collections (Ubi et al., 2022). Insertion presence is therefore a property of a cultivar, not of the species.

In an allopolyploid, every gene is normally present in three homoeologous copies, and those copies are not expressed equally. Surveying the Chinese Spring transcriptome across tissues and developmental stages, Ramirez-Gonzalez et al. (2018) classified homoeologous triads as balanced, as showing dominance of a single homoeolog, or as showing suppression of a single homoeolog, and found that a majority are balanced while a substantial minority depart from equality in a tissue-dependent manner. The same framework extends to groups in which one subgenome copy has been lost (dyads) or duplicated (tetrads). Whether this imbalance is heritable, cultivar-level variation, or a property of the species was only recently addressed, when de novo annotation and differential expression analysis of nine sequenced cultivars across six tissues revealed widespread changes in homoeolog expression bias among cultivars alongside a large conserved core (White et al., 2025). Homoeolog expression bias is thus itself polymorphic, which raises the question of what varies with it.

Transposable elements are an obvious candidate, and the general principle is established outside wheat. Elements inserted near or within genes attract repressive chromatin marks that can spread into adjacent sequence, imposing a cost on expression that scales with proximity (Hollister and Gaut, 2009); they supply stress-responsive regulatory sequence (Makarevitch et al., 2015); and their effects on neighboring transcription are now well cataloged across plant genomes (Hirsch and Springer, 2017; Lanciano and Cristofari, 2020). Where whole populations have been surveyed, insertion polymorphism accounts for a measurable share of expression and phenotypic variation, as shown for the Arabidopsis thaliana mobilome (Quadrana et al., 2016; Stuart et al., 2016) and for tomato (Domínguez et al., 2020). In wheat, the question has been addressed genome-wide within a single genotype: scoring elements lying entirely within gene bodies in Chinese Spring, Bariah et al. (2023) found roughly 36% of genes to carry such an element and reported statistically detectable associations with homoeolog expression category, with elements in exons and untranslated regions implicated more strongly than those in introns. Those associations were, however, quantitatively modest, and a single-genotype design cannot distinguish a genuine effect of insertion from the many other features that differ between the three subgenomes. The problem and the opportunity created by the new pan-genome resources have been set out recently (Gribun and Kashkush, 2026): if insertion is polymorphic between cultivars and expression bias is polymorphic between cultivars, the two can be compared directly, and the comparison is internally controlled in a way that no single-genome analysis can be. Two opposite predictions follow, and this study tests both side by side. Under the suppression prediction, the homoeolog that carries a gene-body element is the one expressed least within its group; this is the direction reported for Chinese Spring (Bariah et al., 2023). Under the dominance prediction, that same homoeolog is the one expressed most; this is the direction favoured when the wheat pan-transcriptome is analysed with the per-observation metric used in its original report (White et al., 2025). Because a single homoeologous group can satisfy both rules at once, the two must be tested against each other rather than one at a time.

In this study, we sequenced young-leaf transcriptomes of six bread wheat cultivars of diverse geographic origins, quantified them against IWGSC RefSeq v2.1, and combined them with transposable element annotations generated independently for each cultivar’s genome assembly, so that insertions were scored in the genome they actually occupy. Our aims were to establish how much of the apparent relationship between gene-body insertion and homoeolog expression bias survives an explicit null model; to ask whether the relationship resides in the groups whose insertion pattern differs between cultivars rather than in insertion presence as such; to determine which element positions and superfamilies carry it; and to test whether it is reproduced in an independently generated dataset. We find that insertion polymorphism, rather than insertion presence, is where the signal lies; that it acts in both directions but far more strongly towards suppression of the inserted homoeolog; that it is carried by intronic rather than coding-sequence insertions, and that it is recovered in all six tissues of the wheat pan-transcriptome once the same statistical treatment is applied.

## Results

### Transcriptome quantification and sample consistency in the six bread wheat cultivars

Data from 18 RNA-Seq libraries (3 biological replicates of 6 bread wheat cultivars) yielded 1,075.3 million read pairs (Table S1). Pseudoalignment rates against the Chinese Spring RefSeq v2.1 transcriptome ranged from 86.61% in Chinese Spring itself to 79.67% in LongReach Lancer and were ordered by genetic distance from the reference, with Norin 61 (84.49%), Jagger (82.20%), CDC Stanley (81.74%) and Mace (81.50%) falling in between (Figure 1a). The 6.9-percentage-point spread highlights cultivar-dependent differences in transcriptome representation relative to the Chinese Spring reference and is why the analyses below are based on comparisons within cultivars rather than on absolute expression levels between them. Principal component analysis of log2-transformed expression cleanly separated the cultivars, with all three biological replicates clustering together and the first three components capturing 23.8%, 16.5%, and 13.4% of the variance (Figure 1b). Pairwise correlations between libraries and hierarchical clustering of the 500 most variable genes yielded the same grouping (Figures S1 and S2).

**Figure 1.**
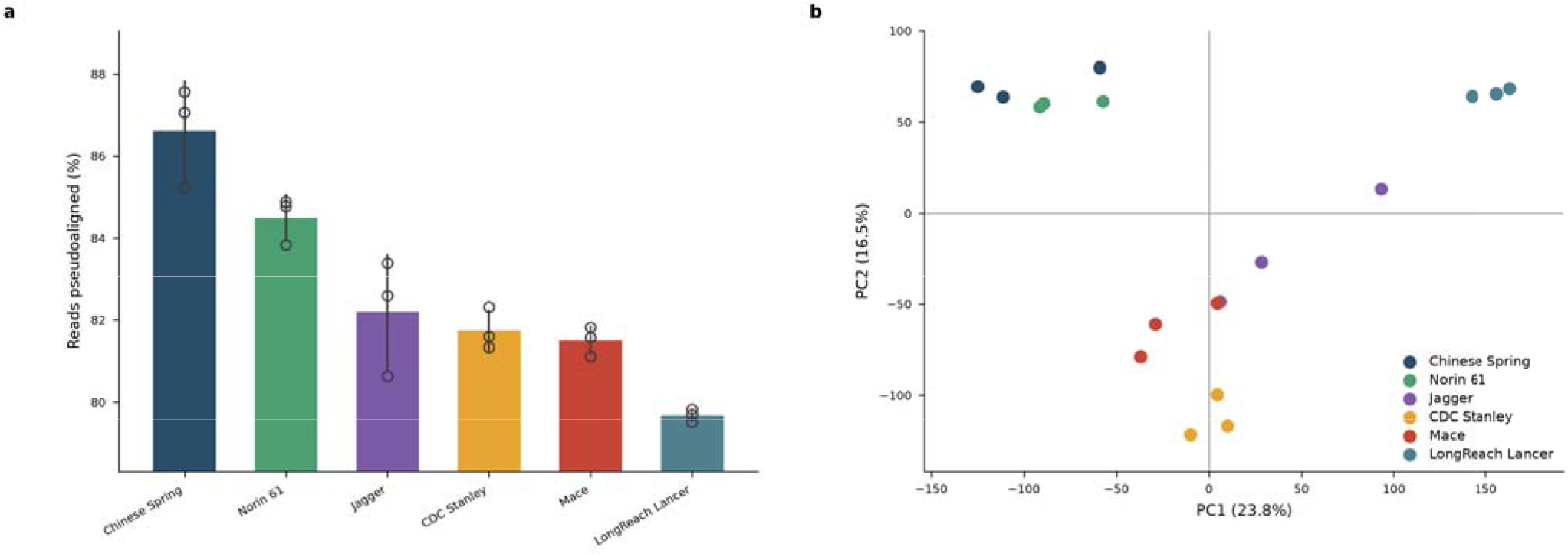
Transcriptome resource. (a) Percentage of read pairs pseudoaligned to the Chinese Spring RefSeq v2.1 transcriptome, by cultivar. Bars show the mean of three biological replicates, whiskers show the standard deviation, and open circles represent the individual libraries. Rates decline with genetic distance from the reference, from 86.61% in Chinese Spring to 79.67% in LongReach Lancer, a 6.9 percentage-point spread. (b) Principal component analysis of log2(TPM+1) across all 266,752 quantified loci and 18 libraries. Replicates are grouped by cultivar on the first two components, which together capture 40.3% of the variance. Cultivars: CS, Chinese Spring; MA, Mace; NO, Norin 61; ST, CDC Stanley; JA, Jagger; LRL, LongReach Lancer.

### Homoeolog expression categories vary between cultivars

Of 25,422 homoeologous groups, 12,239 triads, 2,177 dyads and 328 tetrads were expressed in all six cultivars (Figure 2a). Between 12,981 and 13,467 triads were expressed in any individual cultivar. Assigning each expressed triad to one of seven categories by its nearest ideal expression vector, the balanced class predominated everywhere, but its share varied considerably between cultivars, from 69.3% in Chinese Spring down to 59.4% in LongReach Lancer (Figure 2b). The departures were not symmetrical across the subgenomes. The suppression of the B homoeolog was the most common non-balanced category observed in every cultivar, with occurrences ranging from 965 triads in Chinese Spring to 2,088 in LongReach Lancer. This represents a more than twofold difference between two cultivars of the same species (Table S2a). Homoeolog expression bias in these cultivars is therefore substantial and cultivar-specific.

**Figure 2.**
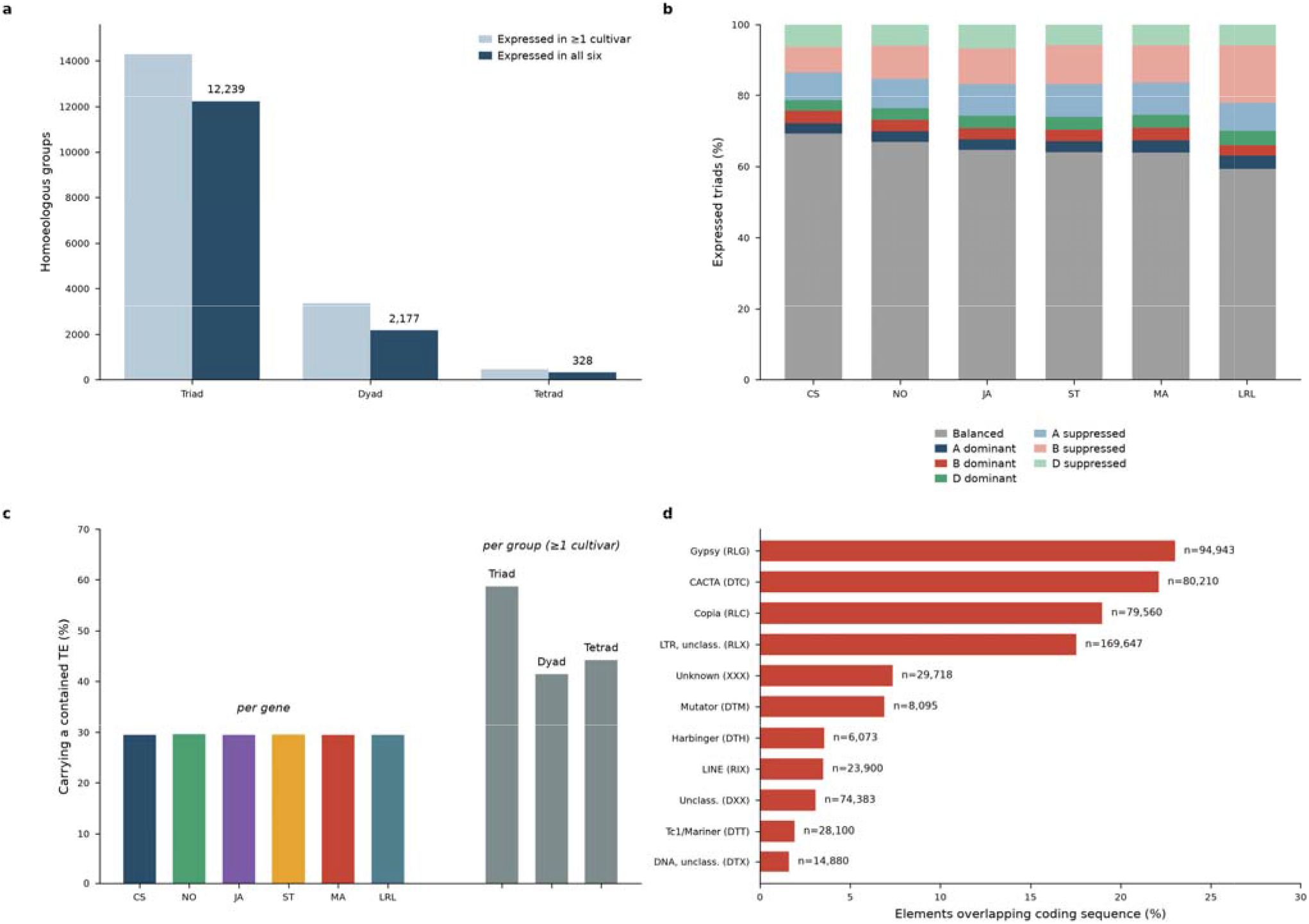
Expressed homoeologous groups, expression categories, and transposable element prevalence and position. (a) Groups expressed in at least one cultivar and in all six, by group type. (b) Distribution of the seven homoeolog expression categories among expressed triads. The balanced class predominates in every cultivar, but its share varies from 69.3% in Chinese Spring to 59.4% in LongReach Lancer. (c) Percentage of genes carrying at least one element lying entirely within the gene body, by cultivar (left), and percentage of homoeologous groups in which at least one cultivar carries such an element, by group type (right). Gene-level prevalence is nearly invariant across six independently assembled genomes, spanning 29.47% to 29.64%. (d) Percentage of contained elements that overlap coding sequence, by superfamily. Overall, 85% of contained elements are intronic and do not touch an exon, but superfamilies differ widely, from Gypsy at 23.0% to unclassified DNA transposons at 1.6%. Cultivars: CS, Chinese Spring; MA, Mace; NO, Norin 61; ST, CDC Stanley; JA, Jagger; LRL, LongReach Lancer.

Across the 12,239 triads expressed in all six cultivars, the category assignment was identical in all six for 6,500 groups and differed in at least one cultivar for the remaining 5,739, so that for 46.9% of triads the pattern of homoeolog expression depends on which cultivar is examined. The stable set is almost entirely balanced: 5,535 of its 6,500 groups are balanced in every cultivar, and only 965 groups, 7.9% of all triads expressed in all six, hold the same non-balanced category throughout the panel. Among the 5,739 variable groups, 4,875 are balanced in at least one cultivar and biased in another, while 864 switch between two different non-balanced categories. LongReach Lancer was the only cultivar that deviated significantly more than any of the others, appearing in 1,032 groups. In contrast, the other five cultivars were represented in only 270 to 313 groups each. This trend corresponds to LongReach Lancer’s position as the cultivar most distant from the reference in terms of pseudoalignment rate. This suggests that part of the excess deviation may be due to the quantification process using Chinese Spring as a reference, rather than being solely a biological difference.

The same contrast appears at the level of individual genes. Testing the 95,622 loci with sufficient coverage against Chinese Spring (Wald test, |log2 fold change| ≥ 1, Benjamini-Hochberg FDR < 0.05), 9,123 genes were differentially expressed in Norin 61, 14,131 in Mace, 16,418 in Jagger, 16,506 in CDC Stanley and 20,654 in LongReach Lancer, and 38,091 genes, 39.8% of those tested, were differentially expressed in at least one cultivar (Table S2b). The fewest and the most were found in the cultivars with the highest and the lowest pseudoalignment rate, respectively, so these counts are an upper bound on genuine expression divergence. Genes belonging to category-variable triads were nevertheless far more likely to be differentially expressed than genes of stable triads, at 43.4% against 13.9% (odds ratio 4.77), so the category switches reflect quantitative differences in expression rather than reassignment of groups lying close to a category boundary.

The triads whose categories vary across cultivars are not a random sample of the transcriptome. Testing the 17,217 genes they contain against the 36,717 genes of all expressed triads, 136 of 614 Gene Ontology terms were significant at a false discovery rate of 5%: 65 enriched and 71 depleted (Table S2c). Enrichment was concentrated in metabolism and transport: hydrolase activity acting on O-glycosyl compounds (1.60-fold, q = 8.0 × 10 ²¹), carbohydrate metabolic process (1.42-fold, q = 5.4 × 10 ¹), transferase activity transferring hexosyl groups (1.57-fold), potassium ion transmembrane transport (1.73-fold), transporter activity (1.39-fold), oxidoreductase activity (1.23-fold) and heme binding (1.30-fold). The depleted terms are as informative: structural constituent of ribosome (0.75-fold), translation (0.75-fold), protein folding (0.64-fold) and intracellular protein transport (0.66-fold) were all under-represented. Cultivar-dependent homoeolog bias therefore falls disproportionately on genes of carbohydrate metabolism, membrane transport and oxidation-reduction, and largely spares the core translational and protein-folding machinery, where balanced expression of all three homoeologs is maintained across the panel.

### Gene-body TEs and their association with homoeolog expression bias

Between 29.47% and 29.64% of genes contained at least one element that lies entirely within the annotated gene body, with each cultivar having between 99,620 and 106,583 such elements (Table 1). It’s noteworthy that this percentage varies by less than 0.2 percentage points across six independently assembled and annotated genomes. When examining homoeologous groups, at least one cultivar contained an element in 58.7% of triads, 41.4% of dyads, and 44.2% of tetrads (Figure 2c and Table S3). In our classification of each element by position, we found that 84.87% to 85.37% were located within introns and did not overlap with any coding exons across all cultivars. The different superfamilies showed significant variation in this aspect. For instance, the Gypsy superfamily had 23.0% of its elements overlapping with coding sequences, followed closely by CACTA at 22.1% and Copia at 19.0%. In contrast, the Mutator superfamily had only 6.9%, and the Tc1/Mariner and unclassified DNA transposons had even lower percentages at 1.9% and 1.6%, respectively (Figure 2d and Table S4).

**Table 1.**
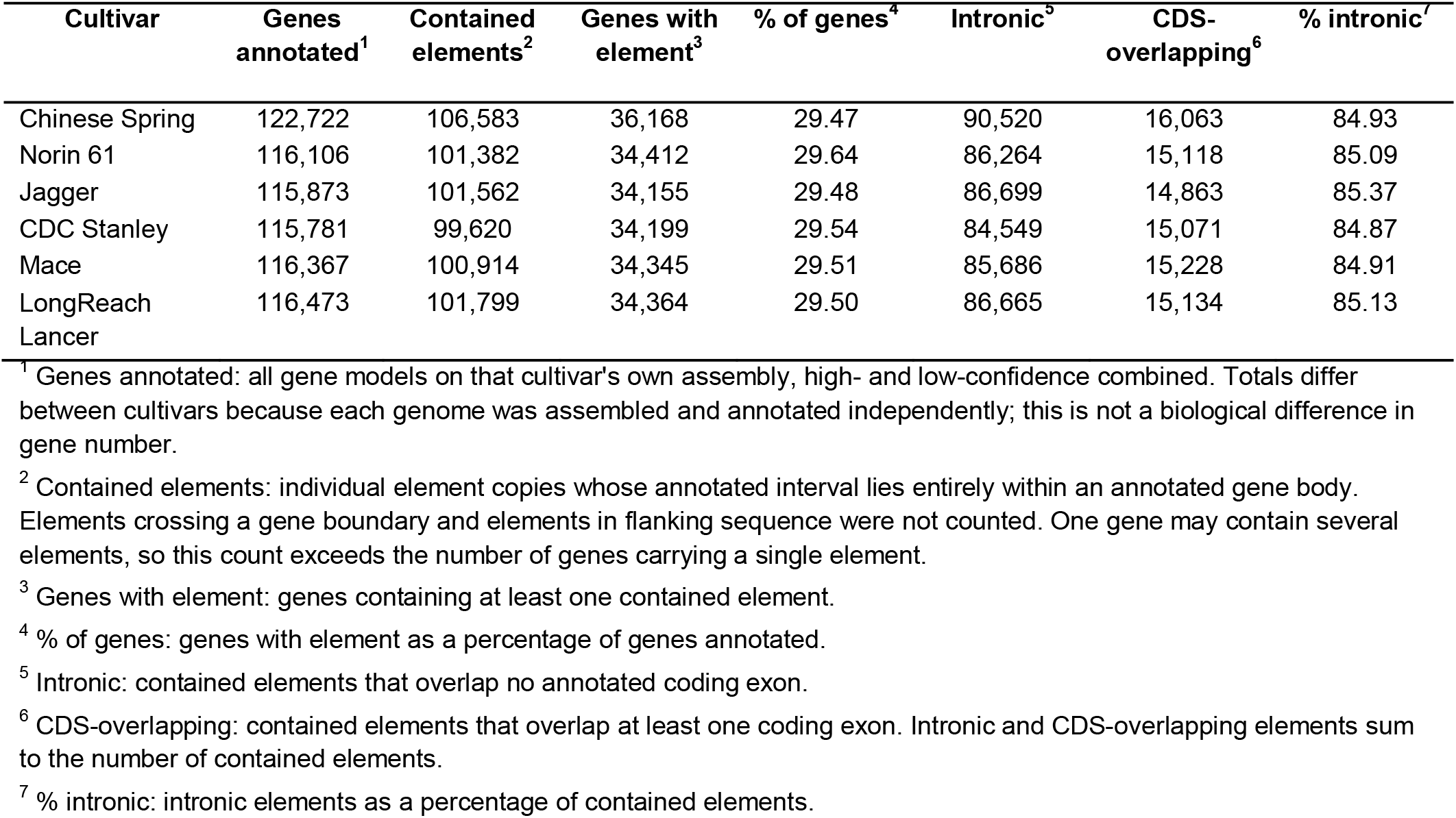
Transposable element prevalence and position within gene bodies, by cultivar.

### Expression bias tracks insertion only where insertion differs between cultivars

Among the 12,887 analyzed groups, 4,648 carried no contained element in any cultivar, 6,547 carried the identical insertion pattern in all six, and only 1,692 differed between cultivars. The first two classes cannot be informative in a comparison between cultivars: where the insertion pattern does not vary, any agreement with expression category is determined by the distribution of categories alone. We therefore tested the full set and the polymorphic subset separately against a permutation null in which insertion-pattern vectors were shuffled 2,000 times within strata defined by group type and present-subgenome combination, with expression categories held fixed (Table 2).

**Table 2.** Support for the suppression and dominance predictions against a permutation null.

| Prediction | Stratum<sup>1</sup> | Observed<sup>2</sup> | Expected ± SD<sup>3</sup> | Fold<sup>4</sup> | Excess<sup>5</sup> | Attributable<sup>6</sup> | p<sup>7</sup> | q<sup>8</sup> |
|---|---|---|---|---|---|---|---|---|
| ***Partitioned by insertion polymorphism*** |  |  |  |  |  |  |  |  |
| Suppression | All analysed groups | 225 | 146.8 ± 11.3 | 1.53 | +78 | 35% | < 0.001 | — |
| Suppression | Polymorphic for insertion | 73 | 10.7 ± 3.2 | 6.83 | +62 | 85% | < 0.001 | — |
| Dominance | All analysed groups | 203 | 184.6 ± 12.5 | 1.10 | +18 | 9% | 0.084 | — |
| Dominance | Polymorphic for insertion | 42 | 16.6 ± 4.0 | 2.53 | +25 | 60% | < 0.001 | — |
| Either direction | All analysed groups | 428 | 331.4 ± 15.7 | 1.29 | +97 | 23% | < 0.001 | — |
| Either direction | Polymorphic for insertion | 115 | 27.3 ± 5.0 | 4.21 | +88 | 76% | < 0.001 | — |
| ***Partitioned by group type*** |  |  |  |  |  |  |  |  |
| Suppression | Triads | 183 | 115.4 ± 10.1 | 1.59 | +68 | 37% | 0.001 | 0.006 |
| Dominance | Triads | 171 | 154.3 ± 11.4 | 1.11 | +17 | 10% | 0.154 | 0.308 |
| Suppression | Dyads | 37 | 25.5 ± 4.7 | 1.45 | +12 | 31% | 0.023 | 0.069 |
| Dominance | Dyads | 28 | 25.1 ± 4.5 | 1.11 | +3 | 10% | 0.577 | 0.865 |
| Suppression | Tetrads | 5 | 5.7 ± 2.2 | 0.88 | -1 | -14% | 0.976 | 0.976 |
| Dominance | Tetrads | 4 | 5.3 ± 2.1 | 0.76 | -1 | -32% | 0.746 | 0.895 |

<sup>1</sup> Stratum: in the upper block, groups are split by whether the insertion pattern differs between the six cultivars (polymorphic) or is identical in all six. In the lower block, all analyzed groups are split by group type without that restriction.
<sup>2</sup> Observed: groups whose expression category matches the prediction in every cultivar. Groups that satisfy both predictions trivially, because their insertion pattern is symmetric and their expression balanced, are excluded from both counts.
<sup>3</sup> Expected ± SD: mean and standard deviation of the observed count across the 2,000 permutations.
<sup>4</sup> Fold: observed divided by expected.
<sup>5</sup> Excess: observed minus expected, rounded to the nearest group.
<sup>6</sup> Attributable: excess as a percentage of the observed count, that is, the fraction of listed groups not explained by chance.
<sup>7</sup> p: upper-tail empirical probability from the permutation distribution. The smallest attainable value is 1/2001, so entries shown as < 0.001 are at that floor and are not exact values.
<sup>8</sup> q: Benjamini–Hochberg correction across the six group-type tests of the lower block. The upper block was not corrected for multiple testing, and is marked —.

Across all analyzed groups, the suppression prediction was supported by 225 groups against 146.8 expected, a 1.53-fold enrichment (p < 0.001), and the dominance prediction by 203 against 184.6 expected, a 1.10-fold enrichment that is not distinguishable from chance (p = 0.084). Restricted to the 1,692 polymorphic groups, the picture changed substantially: 73 groups supported suppression against 10.7 expected, a 6.83-fold enrichment (p < 0.001), and 42 supported dominance against 16.6 expected, 2.53-fold (p < 0.001) (Figure 3a). Taking the two directions together, 115 polymorphic groups showed expression bias tracking the insertion against 27.3 expected, a 4.21-fold enrichment, of which 76% is attributable to the association rather than to chance. Among supporting groups overall, 32.4% were polymorphic for insertion, compared with 7.3% expected. The association between gene-body insertion and homoeolog expression bias therefore resides almost entirely in the groups whose insertion patterns differ between cultivars, and it operates in both directions, but far more strongly towards suppression of the inserted homoeolog.

**Figure 3.**
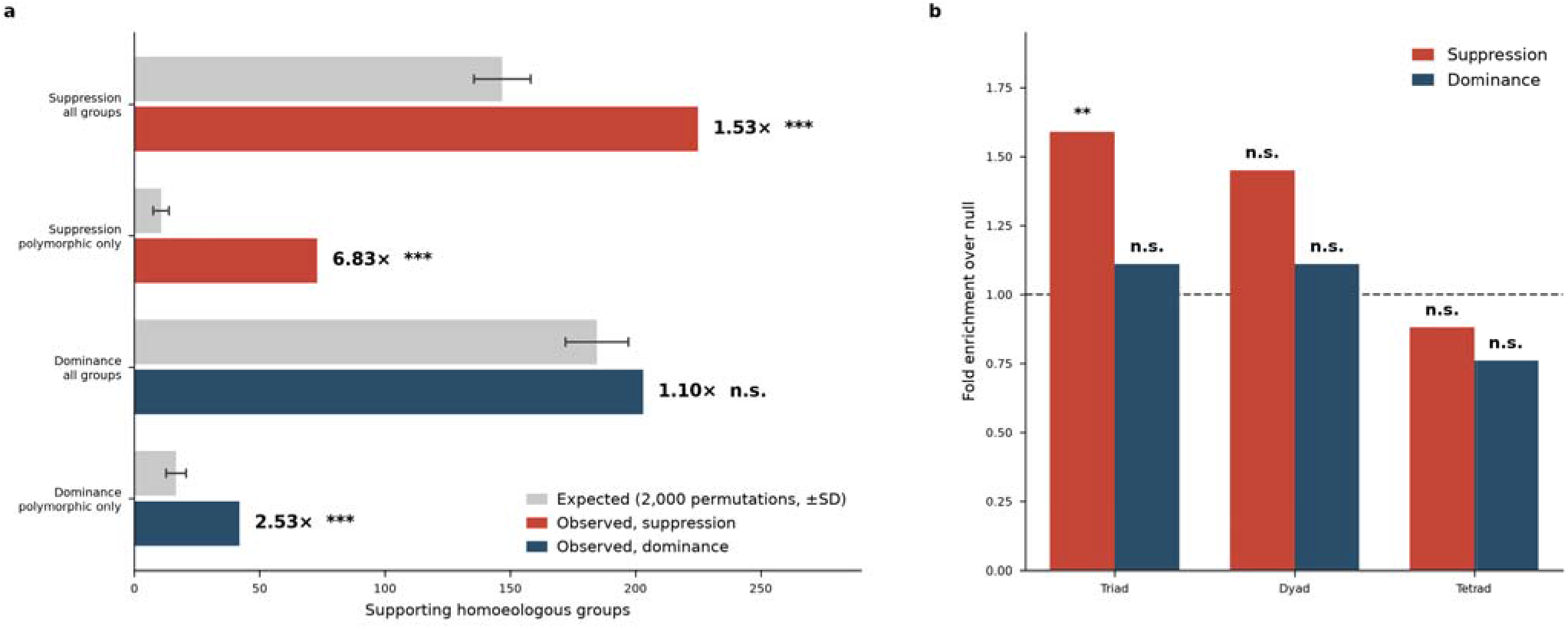
Transposable element insertion and homoeolog expression bias, tested against a permutation null. (a) Observed number of homoeologous groups supporting each prediction (coloured bars) against the number expected when insertion patterns are permuted within strata of group type and present-subgenome combination (grey bars, mean of 2,000 permutations; whiskers give the standard deviation). Fold enrichment is given to the right of each observed bar. Across all analyzed groups, the suppression prediction is supported 1.53-fold more often than chance; restricted to the 1,692 groups whose insertion pattern differs between cultivars, the enrichment rises to 6.80-fold, with 73 groups observed against 10.7 expected. The dominance prediction behaves similarly at lower amplitudes, rising from 1.10-fold across all groups, which is indistinguishable from chance, to 2.53-fold among polymorphic groups. *** p < 0.001; n.s., not significant. (b) Fold enrichment by homoeologous group type, across all analyzed groups. Tetrads contribute nothing to either direction. ** q < 0.01; n.s., not significant.

Partitioned by group type without the polymorphism restriction, suppression was enriched 1.59-fold in triads (q = 0.006) and 1.45-fold in dyads (q = 0.069), while tetrads showed no enrichment in either direction (0.88-fold and 0.76-fold) (Figure 3b). Dominance reached only 1.11-fold in triads (q = 0.308). The effect is thus a triad and dyad phenomenon and should not be described as a general property of homoeologous groups.

If the association arose because elements interrupt coding sequence, it should be carried by the CDS-overlapping minority of insertions. It is not. Rebuilding insertion patterns from intronic elements alone gave 178 suppression-supporting triads against 103.0 expected, a 1.73-fold enrichment (q = 0.004) that is stronger than for all elements combined, whereas patterns built from CDS-overlapping elements alone gave 118 against 84.0 expected, 1.41-fold (q = 0.004) (Figure 4a). For the dominance prediction, intronic elements yielded 135 supporting triads against an expected 135.3, an enrichment of exactly 1.00 (q = 1.000). This is the opposite of the emphasis reported for Chinese Spring alone, where insertions in exons and untranslated regions were implicated more strongly than intronic insertions (Bariah et al., 2023).

**Figure 4.**
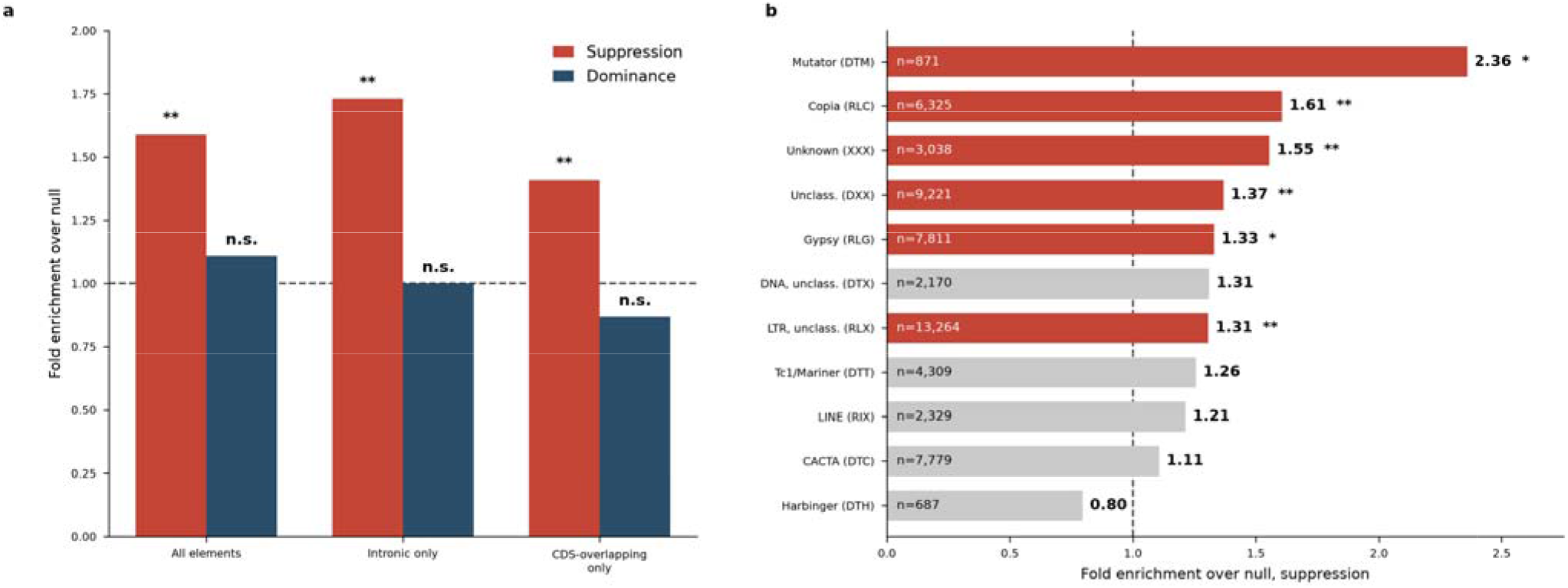
Element position and superfamily. (a) Fold enrichment for triads when insertion patterns are rebuilt from elements in defined positions. If the association arose because elements interrupt coding sequence, it should be carried by the CDS-overlapping minority; instead, intronic elements alone give a stronger enrichment for suppression (1.73-fold) than all elements combined (1.59-fold) or CDS-overlapping elements alone (1.41-fold), while dominance falls to exactly 1.00 for intronic elements. (b) Fold enrichment for suppression in triads, by transposable element superfamily, for the eleven superfamilies present in at least 300 genes. Red bars are significant after Benjamini-Hochberg correction; grey bars are not; the number of genes carrying each superfamily is given inside the bar. Mutator shows the largest effect, and CACTA none at all despite being present in 7,779 genes. ** q < 0.01; * q < 0.05.

Testing each superfamily separately in triads, suppression was significantly enriched for six of the eleven superfamilies present in at least 300 genes: Mutator (2.36-fold, q = 0.020, from 15 supporting groups), Copia (1.61-fold, q = 0.005), unknown elements (1.55-fold, q = 0.005), unclassified elements (1.37-fold, q = 0.005), Gypsy (1.33-fold, q = 0.027) and unclassified LTR elements (1.31-fold, q = 0.005) (Figure 4b). CACTA showed no enrichment at all (1.11-fold, q = 0.432) despite being present in 7,779 genes, which is more than Copia or Gypsy, and it is not explained by position: CACTA elements overlap coding sequence about as often as Gypsy elements do. For the dominance prediction, only Copia departed from expectation, and in the opposite direction, being depleted 0.73-fold (q = 0.033). Full per-superfamily statistics for all group types and both predictions are given in Table S5. Four of the six superfamilies associated with suppression here, Copia, Mutator, unclassified LTR and unknown elements, were also reported as associated with suppression in Chinese Spring alone (Bariah et al., 2023).

#### Replication in the wheat pan-transcriptome

We next asked whether an independently generated dataset gives the same answer. Applying the identical support rule and permutation null to the 12,571 homoeologous groups shared by nine sequenced cultivars across six tissues (White et al., 2025), the suppression prediction was enriched in every tissue, from 1.01-fold in flag leaf to 1.37-fold in grain, with grain surviving correction for multiple testing (q = 0.024) (Figure 5a). Gene-body insertion prevalence in these cultivars was comparable to our own, at 32.9% to 33.4% of all annotated genes and 47.6% to 48.2% of genes belonging to complete homoeologous groups (Figure S3). Analyzed instead with the per-observation chi-square metric used in the original report, the two directions appear comparable in magnitude, an impression that arises because the same group contributes an observation in every cultivar and tissue (Figure S4). The direction is therefore reproduced in a dataset generated in a different laboratory, from different plants, with independently constructed gene models, and in five tissues that were never sampled in our own experiment. Because these cultivars were sequenced to the chromosome level, the same reads could be quantified either against Chinese Spring or against each cultivar’s own genome, allowing the association to be measured independently of the choice of reference. Bias-class assignments agreed between the two alignments for 90.2% of cultivar-by-tissue observations. Where they disagreed, the difference was systematic: alignment to Chinese Spring assigned more observations to the suppressed class, raising it from 18.60% to 20.94% while leaving the dominant class unchanged, and the difference was larger for groups carrying a transposable element (2.49 percentage points) than for element-free groups (1.97 percentage points; Wilcoxon p = 0.00094). Own-genome quantification therefore provides the more conservative estimate of the association. Mean suppression enrichment across tissues was 1.507 under Chinese Spring alignment and 1.239 under own-genome alignment, and the mean absolute excess per tissue was 30.1 and 12.8 groups respectively (Figure 5b). Dominance was essentially unaffected, at 1.12 to 1.19 against 1.10 to 1.16. The association is recovered under both alignments, and grain remained significant under own-genome alignment (1.417-fold, q = 0.005). Per-tissue statistics under both alignments are given in Table S6 and the full agreement matrix in Table S7.

**Figure 5.**
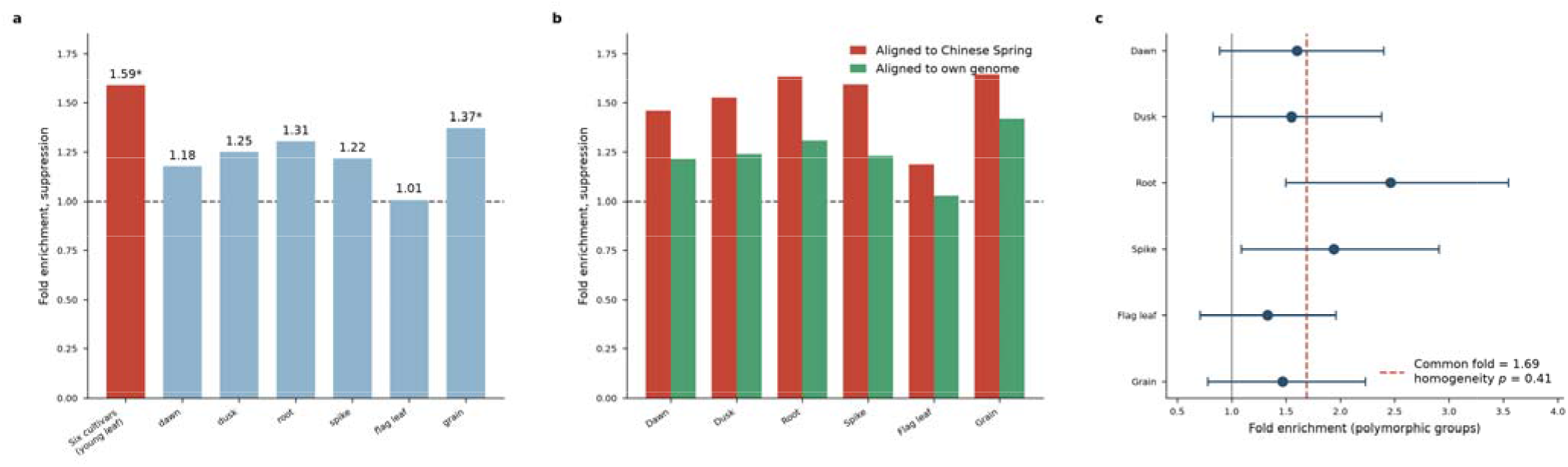
Replication in the wheat pan-transcriptome. (a) Fold enrichment for the suppression prediction in the six-cultivar panel (red) and in each of six tissues of the wheat pan-transcriptome of White et al. (2025) (blue), analyzed with the identical support rule and permutation null. The direction is reproduced in every tissue. * q < 0.05. (b) The same pan-transcriptome groups analyzed from reads aligned to Chinese Spring and from reads aligned to each cultivar’s own genome. The enrichment is recovered under both alignments in every tissue, averaging 1.507 under Chinese Spring alignment and 1.239 under own-genome alignment. (c) Fold enrichment among polymorphic groups by tissue, with bootstrap 95% confidence intervals from 2,000 resamples of groups. All six intervals overlap a common enrichment of 1.69, and the six-tissue homogeneity test is not significant (p = 0.41), so these data provide no evidence that the association differs between tissues.

Enrichment did not differ detectably between the six tissues. Bootstrap confidence intervals on the per-tissue folds all overlapped a common enrichment of 1.69 among polymorphic groups, and a six-tissue homogeneity test was not significant (p = 0.41 for polymorphic groups, p = 0.49 across all groups) (Figure 5c). With approximately 17 polymorphic supporting groups per tissue, the study has little power to detect modest tissue dependence, so this is an absence of evidence rather than evidence of absence; it does, however, mean that these data provide no support for the association being specific to any one tissue or developmental stage.

#### Candidate loci

Selecting groups whose insertion pattern differs between cultivars and whose expression matches the suppression prediction in every cultivar, excluding the shared balanced class, yields 73 groups comprising 205 genes, of which 74 are the homoeologs whose insertion status varies (Table 3). The corresponding dominance selection yields 42 groups and 42 candidate homoeologs. Because roughly 15% of the suppression list and 40% of the dominance list are expected by chance, these are candidates for validation rather than established cases, and the lists cannot indicate which individual members are real. The complete lists are given in Tables S8 and S9, and the corresponding pan-transcriptome candidates in Table S10.

**Table 3.** Twenty highest-priority candidate loci at which a polymorphic insertion coincides with suppression of the inserted homoeolog in every cultivar.

| No. | Gene identifier<sup>1</sup> | Chr | Sub<sup>2</sup> | Annotation<sup>3</sup> | TE present in<sup>4</sup> | TE absent in<sup>4</sup> |
|---|---|---|---|---|---|---|
| 1 | TraesCS3B03G0668400 † | 3B | B | Cytochrome P450 | MA, LRL, ST, JA | CS, NO |
| 2 | TraesCS2D03G1091300 † | 2D | D | Cell division protein ftsZ | JA | CS, MA, LRL, ST, NO |
| 3 | TraesCS2A03G0101800 | 2A | A | Cytochrome P450 | MA, ST, JA | CS, LRL, NO |
| 4 | TraesCS5A03G0618300 | 5A | A | Phosphoribosylaminoimidazole-succinocarboxamid | LRL, ST, JA | CS, MA, NO |
| 5 | TraesCS1B03G1078600 | 1B | B | tRNA-2-methylthio-N(6)-dimethylallyladenosine | LRL, ST, JA | CS, MA, NO |
| 6 | TraesCS6B03G1091800 | 6B | B | Serine incorporator | LRL, ST, JA | CS, MA, NO |
| 7 | TraesCS1B03G0874400 | 1B | B | Proteasome subunit alpha type | MA, LRL, NO | CS, ST, JA |
| 8 | TraesCS1B03G0517800 | 1B | B | Coatomer subunit gamma | MA, ST, JA | CS, LRL, NO |
| 9 | TraesCS7B03G0788500 | 7B | B | Receptor-like protein kinase | LRL, ST, NO | CS, MA, JA |
| 10 | TraesCS7B03G1124800 | 7B | B | WRKY transcription factor | MA, ST, JA | CS, LRL, NO |
| 11 | TraesCS6B03G0359800 | 6B | B | Calcium-dependent lipid-binding domain protein | ST, NO | CS, MA, LRL, JA |
| 12 | TraesCS2B03G1456000 | 2B | B | Pentatricopeptide repeat-containing protein | ST, JA | CS, MA, LRL, NO |
| 13 | TraesCS7B03G0807500 | 7B | B | F-box/kelch-repeat protein | LRL, ST | CS, MA, NO, JA |
| 14 | TraesCS3B03G0522200 | 3B | B | Cysteine proteinase inhibitor | MA, LRL, ST, JA | CS, NO |
| 15 | TraesCS7B03G0157100 | 7B | B | zinc ion binding protein | MA, LRL, ST, JA | CS, NO |
| 16 | TraesCS6A03G0042300 | 6A | A | RNA binding | MA, LRL | CS, ST, NO, JA |
| 17 | TraesCS2B03G0935600 | 2B | B | Kelch repeat-containing protein | LRL, ST | CS, MA, NO, JA |
| 18 | TraesCS1B03G0729200 | 1B | B | Rhomboid family protein | MA, JA | CS, LRL, ST, NO |
| 19 | TraesCS5B03G0834800 | 5B | B | WD-repeat protein, putative | ST, JA | CS, MA, LRL, NO |
| 20 | TraesCS2B03G0979500 | 2B | B | Proliferating cell nuclear antigen | ST, JA | CS, MA, LRL, NO |

Selected from the 74 candidate homoeologs by cross-dataset replication first, then by the evenness of the carrier versus non-carrier split, which determines how informative a locus is for experimental validation, and finally by the availability of a functional annotation. The complete list is given in Table S8. These are candidates, not validated loci: roughly 15% of the 74 are expected to arise by chance, and the table cannot indicate which.
<sup>1</sup> Gene identifier: IWGSC RefSeq v2.1. The identifier names the homoeolog whose insertion status varies between cultivars, not the homoeologous group. † marks loci recovered independently in the pan-transcriptome of White et al. (2025).
<sup>2</sup> Sub: the subgenome, A, B or D, carrying the variable insertion.
<sup>3</sup> Annotation: human-readable description from the IWGSC RefSeq v2.1 functional annotation, truncated where the original description exceeds the column width.
<sup>4</sup> Cultivars: CS, Chinese Spring; MA, Mace; LRL, LongReach Lancer; ST, CDC Stanley; NO, Norin 61; JA, Jagger. A cultivar listed under ‘TE present in’ carries at least one element lying entirely within the gene body of the named homoeolog; the remaining cultivars carry none.

Four loci illustrate the patterns directly (Figure 6). At a WRKY transcription factor on chromosome 7B (TraesCS7B03G1124800) a single Copia element occupies the B homoeolog in Mace, CDC Stanley and Jagger, all three of which show the B homoeolog suppressed, while Chinese Spring, Norin 61 and LongReach Lancer carry no element and are balanced. At a cytochrome P450 on 3B (TraesCS3B03G0668400), eight to nine Copia and Gypsy elements occupy the B homoeolog in four cultivars that show B suppressed or D dominant, against two element-free and balanced cultivars; this group is one of two also recovered independently in the pan-transcriptome, where Norin 61 is likewise element-free. Two further loci, a phosphatidylserine decarboxylase proenzyme on 4B and an EamA-like transporter on 2B, show the converse: a Gypsy element in the B homoeolog coinciding with dominance of that homoeolog in four and five cultivars, respectively.

**Figure 6.**
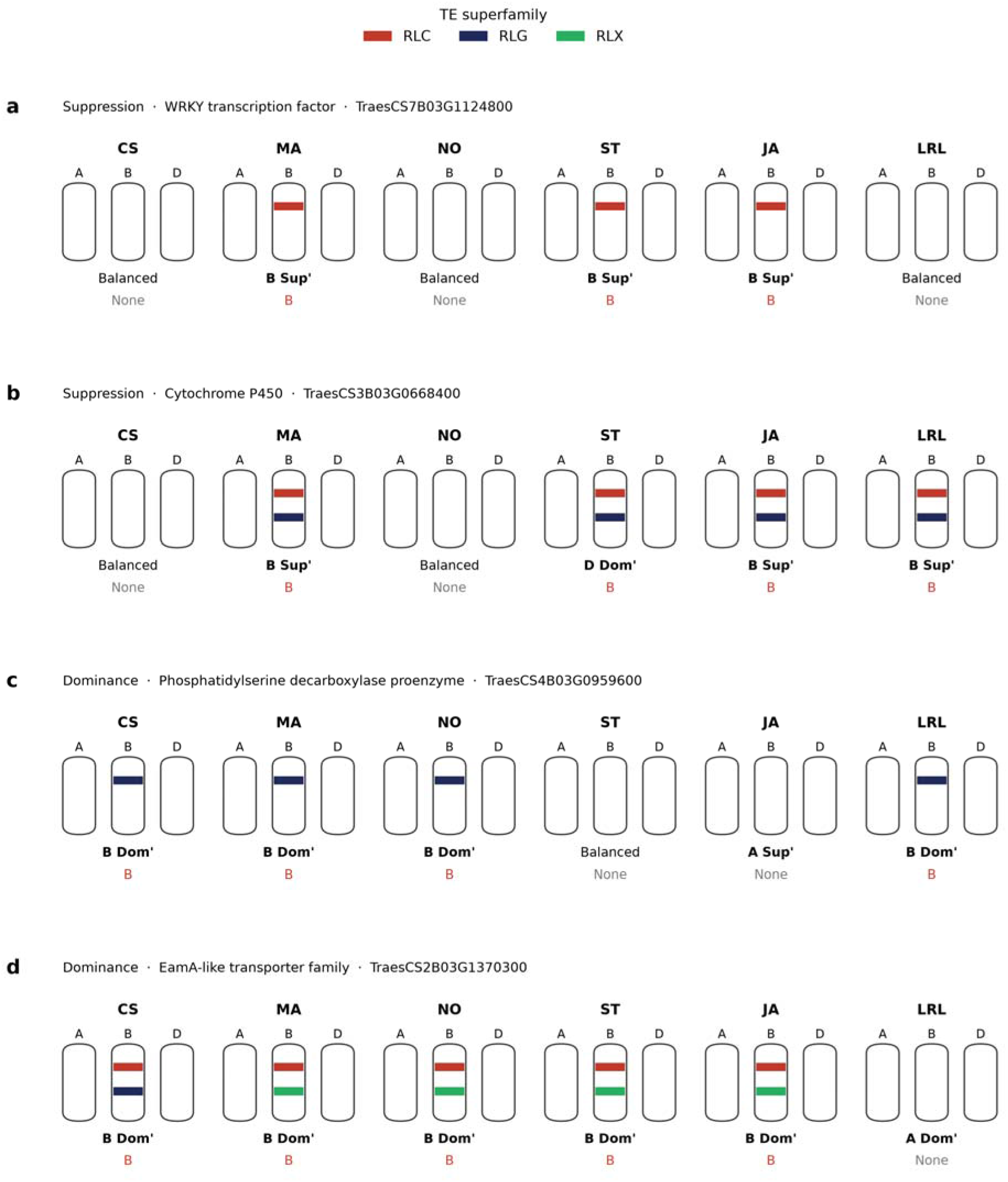
Transposable element insertion patterns and homoeolog expression categories at four homoeologous gene groups. Each row shows one triad across the six cultivars. Within each cultivar the three capsules represent the A, B and D homoeologs; coloured bars mark elements lying entirely within the gene body, coloured by superfamily, with one bar per superfamily present and bar position within the capsule schematic rather than to scale. Beneath each cultivar the first line gives the expression category and the second the insertion pattern, with the inserted subgenome in red. Gene identifiers name the B homoeolog, which carries the polymorphic insertion in all four examples. (a) WRKY transcription factor: a single Copia element in Mace, CDC Stanley and Jagger, all of which show the B homoeolog suppressed, against three element-free and balanced cultivars. (b) Cytochrome P450: eight to nine Copia and Gypsy elements in four cultivars; this group is also recovered independently in the pan-transcriptome. (c) Phosphatidylserine decarboxylase proenzyme: a single Gypsy element in four cultivars, all B dominant. (d) EamA-like transporter: two elements in five cultivars, all B dominant, against an element-free LongReach Lancer. Cultivars: CS, Chinese Spring; MA, Mace; NO, Norin 61; ST, CDC Stanley; JA, Jagger; LRL, LongReach Lancer. Only elements lying entirely within the gene body were counted, so insertions in promoters or flanking sequence are not shown.

The candidate loci do not cluster within any specific functional class. Among the 299 candidates identified across both directions and datasets, only twelve have stress-related annotations, and just one is associated with classical abiotic stress. Additionally, when poorly annotated genes are excluded, the functional composition of the lists for suppression and dominance lists is statistically similar. This suggests that these elements affect the genes they occupy indiscriminately, rather than targeting a coherent pathway. However, at the level of the entire supporting set, a functional trend can be observed: suppression-supporting triads that were monomorphic for both insertion and expression showed a significant enrichment for peroxidase activity (9.0-fold, q = 0.0006), response to oxidative stress (8.4-fold), and heme binding (5.4-fold). In contrast, the corresponding dominance set exhibited broader terms with smaller effect sizes (Figure S5).

## Discussion

Transposable elements occupy gene bodies in roughly three in ten wheat genes, a figure that proved remarkably stable across six independently assembled genomes, and the great majority of those insertions lie in introns. Whether such insertions affect the unequal expression of homoeologous copies has, until now, been tested within single genotypes, where the three subgenomes differ in countless ways besides insertions. By comparing six cultivars, we could ask a more pointed question: does expression bias change where insertions occur? The answer is that it does, and that this is where essentially the whole signal resides. Across all analyzed groups, the suppression prediction exceeded a permutation null of 1.53-fold, an effect so modest that it would reasonably be dismissed; restricted to groups whose insertion patterns differ between cultivars, the same test yielded 6.83-fold. The groups in which insertion does not vary dilute the comparison and constitute 87% of the analyzed set.

The association acts in both directions, but not equally. Taking suppression and dominance together, 115 of the 1,692 polymorphic groups showed expression bias tracking the insertion against 27.3 expected. Separated, suppression of the inserted homoeolog reached 6.83-fold, while dominance reached 2.53-fold; across all analyzed groups, dominance was not distinguishable from chance at all. A directional asymmetry of this kind is what the silencing literature would predict: elements attract repressive chromatin that can spread into flanking sequence (Hollister and Gaut, 2009), which lowers expression of the copy carrying them, whereas raising the relative expression of an inserted homoeolog requires some additional explanation, such as the element supplying regulatory sequence (Makarevitch et al., 2015) or the suppression of a competing homoeolog elsewhere in the group. Our dominance candidates are, in fact, dominated by the latter situation, in which the inserted homoeolog is balanced while its counterpart is suppressed, and should not be read as elements that elevate expression.

Two features of the data argue against the simplest mechanical explanation, that elements disrupt coding sequence. First, insertion patterns reconstructed from intronic elements alone yielded a stronger enrichment (1.73-fold) than those reconstructed from CDS-overlapping elements (1.41-fold), and intronic elements showed no dominance signal whatsoever. Second, CACTA elements showed no association despite occupying 7,779 genes and overlapping coding sequence as often as Gypsy elements do, which rules out position as a sufficient explanation for which superfamilies matter. Both observations point away from coding disruption and towards a chromatin-level effect, although our data cannot demonstrate one. They also differ from the pattern reported for Chinese Spring alone, where exonic and untranslated-region insertions appeared more consequential than intronic insertions (Bariah et al., 2023); the discrepancy may reflect a different unit of comparison, since a single-genotype analysis compares subgenomes, whereas ours compares cultivars.

The pan-transcriptome analysis served two purposes. It replicated the direction of the association in a dataset generated in a different laboratory, from different plants, with independently constructed gene models, and in five tissues we never sampled. Because the same reads could be aligned either to Chinese Spring or to each cultivar’s own genome assembly, it also allowed the association to be measured independently of the choice of reference. Under own-genome alignment the mean enrichment was 1.239 against 1.507 under a common reference, and the mean absolute excess 12.8 against 30.1 groups per tissue. The difference was not specific to transposable elements, since the suppressed class shifted for element-free groups as well, but it was larger for element-carrying groups, and we therefore report the own-genome values as the conservative estimate. The association is recovered under both alignments.

We found no evidence that the association differs between tissues. All six per-tissue estimates were consistent with a single common enrichment, and the homogeneity test was far from significant. This is worth stating because an earlier reading of the same data, contrasting young vegetative tissue against post-anthesis flag leaf, suggested a developmental contrast that formal testing does not support; with roughly 17 polymorphic supporting groups per tissue, the design has little power, and the correct conclusion is that these data neither establish nor exclude tissue-dependence.

Several limitations bound the interpretation. Only elements lying entirely within annotated gene bodies were counted, so insertions in promoters and other flanking sequence, arguably the most likely to affect transcription, are absent from the analysis entirely. Expression categories register a change of category rather than a change of magnitude, so a homoeolog whose expression halves without crossing a category boundary counts as unchanged, which makes the test conservative. Support requires unanimity across all six cultivars, which discards genes suppressed in four. Element annotation depends on the completeness of each assembly, and the PGSB annotation records nested and fragmented elements separately, so element counts exceed the number of historical insertion events. Most importantly, this is an association measured in a single set of plants with no perturbation: the element may affect expression, the element and the expression state may both follow from local chromatin, or the haplotype carrying the element may differ in other ways.

Distinguishing these requires breaking the linkage between element and background, which a segregating population would permit and this design does not.

What the data do support is a reframing of the question. The relevant comparison is not whether a gene carries a transposable element, which a third of wheat genes do, but whether it carries one in some cultivars and not others. In that comparison, the relationship with homoeolog expression bias is reproducible, directionally asymmetric, carried by intronic elements, and about four times stronger than chance after a conservative null. The 74 suppression candidates and 42 dominance candidates mentioned here provide a solid foundation for experiments aimed at determining causation. Among these candidates, the most informative ones are those with an even split between carrier and non-carrier cultivars.

## Material and Methods

### Plant material

Six bread wheat (*Triticum aestivum*, 2n = 6x = 42, AABBDD) cultivars of diverse geographic origin were used: Chinese Spring (China), Mace and LongReach Lancer (Australia), CDC Stanley (Canada), Norin 61 (Japan) and Jagger (United States). All six are among the cultivars assembled to the chromosome level by the 10+ Wheat Genomes Project (Walkowiak et al., 2020), which enabled transposable element annotation to be derived from each cultivar’s own genome rather than inferred from a common reference. Plants were grown in a greenhouse facility at Ben-Gurion University of the Negev under controlled conditions. Leaves were harvested four weeks after germination. Three biological replicates, each an independently grown plant, were maintained per cultivar, giving 18 samples in total.

### RNA extraction, library preparation and sequencing

Total RNA was extracted from young leaves using the Quick-RNA Plant MiniPrep kit (Zymo Research) according to the manufacturer’s instructions, and RNA concentration was determined by NanoDrop spectrophotometry. RNA quality and integrity were assessed using a TapeStation (software version 4.1.1) at the Technion Genomic Center and independently on 1.5% agarose gels, on which clear, unfragmented ribosomal bands were observed; all 18 samples were deemed suitable for library preparation. Libraries were prepared with the NEBNext Ultra II Directional RNA Library Prep Kit using poly(A) selection to deplete ribosomal RNA, pooled, and sequenced on an Illumina NextSeq 2000 instrument with a P3 flow cell over 300 cycles to generate 150 bp paired-end reads. Base-call quality was assessed on the Phred scale, with scores above 30 taken to indicate base-call accuracy exceeding 99.9%.

### Read processing and expression quantification

Raw read quality was assessed with FastQC v0.11. Adaptor sequences and low-quality bases were removed with Trim Galore v0.6.7-1 using default parameters, and post-trimming quality was reconfirmed with FastQC. Transcript abundance was estimated with kallisto v0.52.0 (Bray et al., 2016) using default parameters and an index built from the complete IWGSC RefSeq v2.1 transcriptome of Chinese Spring (Zhu et al., 2021), comprising 295,914 transcripts from 266,752 loci and including both high– and low-confidence gene models; low-confidence models were retained in the index so that reads originating from them would not be misassigned to high-confidence genes. Transcripts per million were summed across transcripts to give gene-level estimates. A gene was considered expressed in a cultivar when its mean TPM across the three biological replicates exceeded 0.5. The sensitivity of downstream counts to this threshold, and to the alternative of requiring every homoeolog of a group to exceed it, is reported in Figure S6. Differential expression between cultivars was tested with PyDESeq2 v0.5.4 on gene-level estimated counts for the 95,622 loci with at least 10 counts in at least three libraries, with cultivar as the only factor and Wald tests of each cultivar against Chinese Spring and Benjamini-Hochberg correction; genes with an adjusted p-value below 0.05 and an absolute log2 fold change of at least 1 were called differentially expressed.

### Homoeologous groups and expression categories

Homoeologous group assignments were taken from the curated set of Ramirez-Gonzalez et al. (2018) and expressed in RefSeq v2.1 gene identifiers. Groups were classified by composition as triads, carrying exactly one copy in each of the A, B and D subgenomes; dyads, in which one subgenome copy is absent; or tetrads, in which one subgenome copy is duplicated. For each group in each cultivar the relative contribution of each homoeolog to the group total was computed from mean TPM, giving a point in the two-dimensional simplex, and assigned to the nearest of seven ideal vectors by Euclidean distance: balanced (1/3, 1/3, 1/3); dominance of each subgenome in turn (1, 0, 0 and its permutations); and suppression of each subgenome in turn (0, 1/2, 1/2 and its permutations). Groups in which any homoeolog fell below the expression threshold were recorded as not expressed in that cultivar and excluded from all category-based analyses. Category assignment was also performed on each replicate separately to quantify replicate agreement (Figure S7).

### TE annotation and insertion patterns

Transposable element annotations produced by PGSB using the Triticeae TE library, and the projected gene models of the same assemblies, were obtained for each of the six cultivars from the 10+ Wheat Genomes Project data repository hosted at IPK Gatersleben (Walkowiak et al., 2020). Because both the element annotation and the gene models are defined on each cultivar’s own chromosome-scale assembly, insertion was scored in the genome of the chromosome the element actually occupies rather than projected onto a common reference. An element was scored as present in a gene only when its annotated interval lay entirely within the annotated gene body; elements overlapping a gene boundary and elements in flanking sequence were excluded, following the definition used by Bariah et al. (2023) so that prevalence figures remain comparable. Each contained element was classified as intronic when it overlapped no annotated coding exon and as CDS-overlapping otherwise, and was assigned to a superfamily using the PGSB three-letter codes, which follow the unified classification of Wicker et al. (2007). Cultivar gene identifiers were linked to Chinese Spring v2.1 identifiers through the correspondence table distributed with the 10+ Wheat Genomes gene projections. For each homoeologous group in each cultivar, an insertion pattern was recorded as the set of subgenomes carrying at least one contained element, giving eight possible states from no insertion to insertion in all three subgenomes. A group was classed as polymorphic for insertion when this pattern was not identical across all six cultivars. As an independent check on the annotation source, the CLARITE annotation distributed with IWGSC RefSeq v2.1 was compared with the PGSB annotation for Chinese Spring across the 104,375 genes that could be matched, yielding 85.6% concordance at the gene-level presence/absence (Figure S8).

### Support rule for the two predictions

A group was scored as supporting the suppression prediction in a given cultivar when its insertion pattern was asymmetric, meaning that at least one but not all present subgenomes carried a contained element, and its expression category was consistent with reduced relative expression of an inserted homoeolog. Where a single subgenome X carried an element, this condition is met when X is the suppressed subgenome or when a non-inserted subgenome is the dominant one. Where two subgenomes carried elements, it is met when the single non-inserted subgenome is the dominant one. The dominance prediction is the exact mirror of these conditions. Symmetric insertion patterns, in which every present subgenome carries an element, or none does, satisfy both predictions whenever expression is balanced; these groups form a shared class that carries no information about which prediction is correct and were excluded from every test reported here. A group was scored as supporting a prediction overall only when every cultivar independently supported it, and the group was expressed in all six cultivars.

### Permutation test and confidence intervals

Significance was assessed by permutation rather than by a parametric test, because the quantity of interest is the pairing of two categorical variables whose marginal distributions are both strongly non-uniform. The six-element vector of insertion patterns belonging to each group was shuffled among groups 2,000 times within strata defined jointly by group type and by which subgenomes are present, while expression categories were held fixed. This preserves the marginal distributions of insertion patterns and expression categories, the correlation of insertion patterns between cultivars within a group, and the composition of each stratum, and asks only whether the observed pairing of pattern with category exceeds what random assignment would produce. Empirical p-values are the proportion of permutations reaching or exceeding the observed count and are floored at 1/2001, so values are reported as p < 0.001 rather than as point estimates; where several related tests were performed, they were corrected by the Benjamini-Hochberg procedure and are reported as q. The identical framework was applied to subsets defined by element position, by superfamily, and by whether the insertion pattern differs between cultivars. Because the same homoeologous groups are tested in every tissue of the pan-transcriptome, bootstrap confidence intervals on per-tissue enrichment were obtained by resampling groups with replacement 2,000 times with a single common resample applied across all tissues, and homogeneity across tissues was tested by a chi-square statistic comparing observed counts with the counts expected under a single common enrichment.

### Pan-transcriptome analysis and reference-independent quantification

Published data from the wheat pan-transcriptome of White et al. (2025) were reanalyzed. That study sampled nine sequenced cultivars (ArinaLrFor, Jagger, Julius, LongReach Lancer, CDC Landmark, Mace, Norin 61, CDC Stanley and SY Mattis) in triplicate across six tissues: whole aerial organs and whole roots harvested separately at the three-leaf stage (Zadoks GS13), with aerial organs sampled both four hours after dawn and at dusk; complete spike at heading (GS59); flag leaf seven days post anthesis (GS71); and whole grain fifteen days post anthesis (GS77). Grain was assayed in seven of the nine cultivars and the remaining tissues in all nine. Homoeologous group assignments, per-tissue homoeolog expression proportions and bias-class calls were taken from the published deposit, restricted to groups with a one-to-one-to-one relationship across all three subgenomes in every line. The support rule and permutation null described above were applied unchanged, requiring every cultivar present in a given tissue to support the prediction. To assess the association independently of the reference used, we took advantage of the two count matrices published by the authors in which the same reads were quantified either against the Chinese Spring reference or against each cultivar’s own genome assembly; bias classes were derived independently from each matrix by the same nearest-ideal-vector rule used for our own data, and the complete analysis was repeated on both. Transposable element annotations for the nine cultivars were obtained and processed exactly as described for the six cultivars.

### Functional annotation and enrichment

Human-readable gene descriptions, Pfam domains, and Gene Ontology terms were taken from the IWGSC RefSeq v2.1 functional annotation. Gene Ontology enrichment was assessed using Fisher’s exact test against the background of all annotated genes in the analyzed set, with Benjamini-Hochberg correction across terms; no semantic reduction of the term list was applied. For triads whose expression category varies between cultivars, the study set comprised the genes of those triads, and the background comprised the genes of all triads expressed in all six cultivars; terms annotated to fewer than 10 background genes were not tested. The broad functional categories used to summarise candidate lists in the text were automatically assigned via ordered keyword matching against Gene Ontology term names and annotation text, and are indicative rather than curated.

### Software, reproducibility and data availability

Analyses were performed in Python 3.11 using pandas (McKinney, 2010), NumPy (Harris et al., 2020), SciPy (Virtanen et al., 2020) and statsmodels (Seabold and Perktold, 2010); differential expression was tested with PyDESeq2 (Muzellec et al., 2023), a Python implementation of the DESeq2 model (Love et al., 2014); figures were produced with Matplotlib (Hunter, 2007). Read quality was assessed with FastQC v0.11 (https://www.bioinformatics.babraham.ac.uk/projects/fastqc/) and adaptor and quality trimming with Trim Galore v0.6.7-1 (https://www.bioinformatics.babraham.ac.uk/projects/trim_galore/). Random seeds were fixed for all permutation and bootstrap procedures, so every figure and every number reported here can be regenerated from the deposited inputs.

## Data availability statement

The wheat pan-transcriptome data reanalyzed here are available from Zenodo (record 16964999) as described by White et al. (2025). Transposable element and gene annotations for the ten wheat genomes are available from the IPK Galaxy server. All tables underlying the figures, the complete candidate-gene lists, the per-gene transposable-element position data, and the gene-level expression matrices are provided as Supporting Information.

Seven data files accompany the manuscript. Data S1 provides the homoeologous group annotations: 18,364 triads, 6,307 dyads, and 751 tetrads, along with their member gene identifiers and chromosomal coordinates, together with an index of all 70,710 member genes. Data S2 provides per-gene records of transposable element insertions: 39,872 genes with presence, element count, and superfamily composition for each of the six cultivars. Data S3 provides the positions of all 198,418 elements, each classified as intronic or CDS-overlapping. Data S4 gives the tables underlying every figure and every numerical statement in the text, including the permutation results under both nulls, the per-position and per-superfamily association statistics, and the pan-transcriptome results under both metrics. Data S5 provides gene-level expression matrices (transcripts per million) and estimated counts for 165,052 genes across the 18 libraries. Data S6 provides the complete Gene Ontology enrichment results for each tested set. Data S7 provides the full differential expression output, log2 fold change, and adjusted p-value for all 95,622 genes tested in each cultivar relative to Chinese Spring.

## Acknowledgements

We would like to thank Vadim Khasdan for his assistance with the greenhouse and lab preparations. We also thank Ibrahim Kashkush, Faculty of Medicine, Al-Quds University, Abu Dis, Jerusalem, Palestine, for his great assistance in the computational analysis. This work was supported by a grant from the Israel Science Foundation (grant# 1311/21) to K. K.

## Supporting information

Figure S1. Verification of cultivar labeling by pairwise library correlation.

Figure S2. Expression heatmap of the 500 most variable genes.

Figure S3. Transposable element prevalence in the wheat pan-transcriptome.

Figure S4. Per-tissue association in the pan-transcriptome under the per-observation chi-square metric.

Figure S5. Gene Ontology enrichment of supporting groups.

Figure S6. Sensitivity of expressed-group counts to the expression threshold.

Figure S7. Replicate agreement in expression-category assignment.

Figure S8. Comparison of PGSB and CLARITE transposable element annotations for Chinese Spring.

Table S1. Plant material, sequencing yield, and pseudoalignment.

Table S2. Homoeolog expression category counts among triads (S2a), genes differentially expressed between each cultivar and Chinese Spring (S2b), and Gene Ontology enrichment of triads whose expression category varies between cultivars (S2c).

Table S3. Transposable element insertion state by homoeologous group type.

Table S4. Position of contained elements by superfamily.

Table S5. Per-superfamily association statistics, all group types and both predictions.

Table S6. Pan-transcriptome association statistics under both alignment strategies.

Table S7. Bias-class agreement between Chinese Spring and own-genome alignment, by cultivar and tissue.

Table S8. Candidate loci, suppression, six cultivars (73 groups, 74 candidate homoeologs).

Table S9. Candidate loci, dominance, six cultivars (42 groups, 42 candidate homoeologs).

Table S10. Candidate loci from the pan-transcriptome, both directions.

Data S1. Homoeologous group annotations.

Data S2. Genes with transposable element insertions.

Data S3. Transposable element position by gene.

Data S4. Statistics underlying every figure and in-text number.

Data S5. Gene-level expression matrices, TPM and counts.

Data S6. Gene Ontology enrichment results.

Data S7. Differential expression of each gene in each cultivar relative to Chinese Spring.

## Author contribution

L.G. generated the plant material, RNA and sequencing data and performed the initial analysis.

I.K. reanalyzed the publicly available pan-genome RNA-Seq data.

K.K. conceived and supervised the study and manuscript preparation.

