## supplemental figures for "Cultivar-specific transposable element insertions are associated with both suppression and dominance of homoeolog expression in bread wheat"

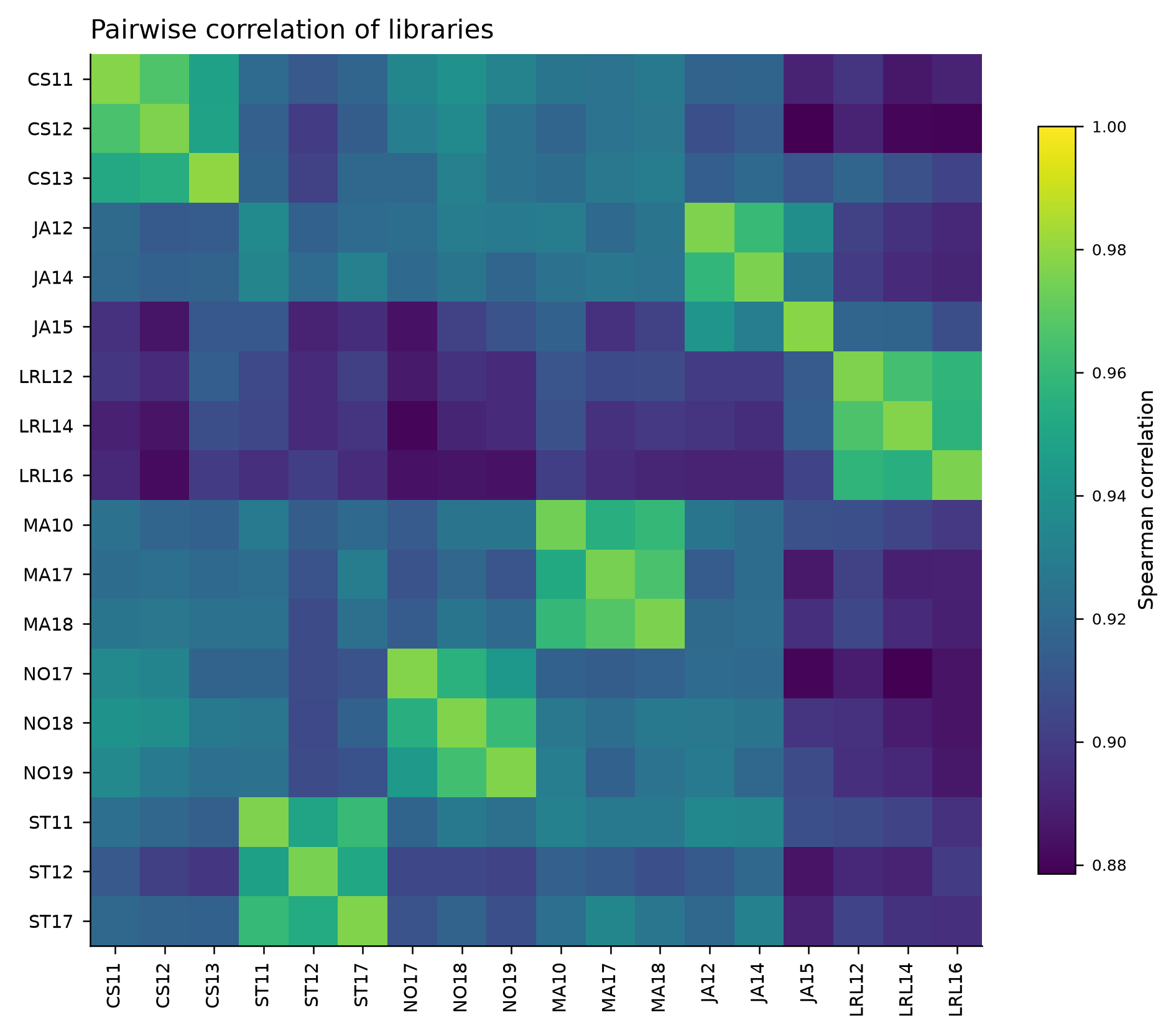

**Figure S1.** Verification of cultivar labeling. Pairwise Spearman correlation between all 18 libraries for gene-level expression. Replicates of the same cultivar form clear diagonal blocks, confirming that library labels are internally consistent and that no sample was misassigned.

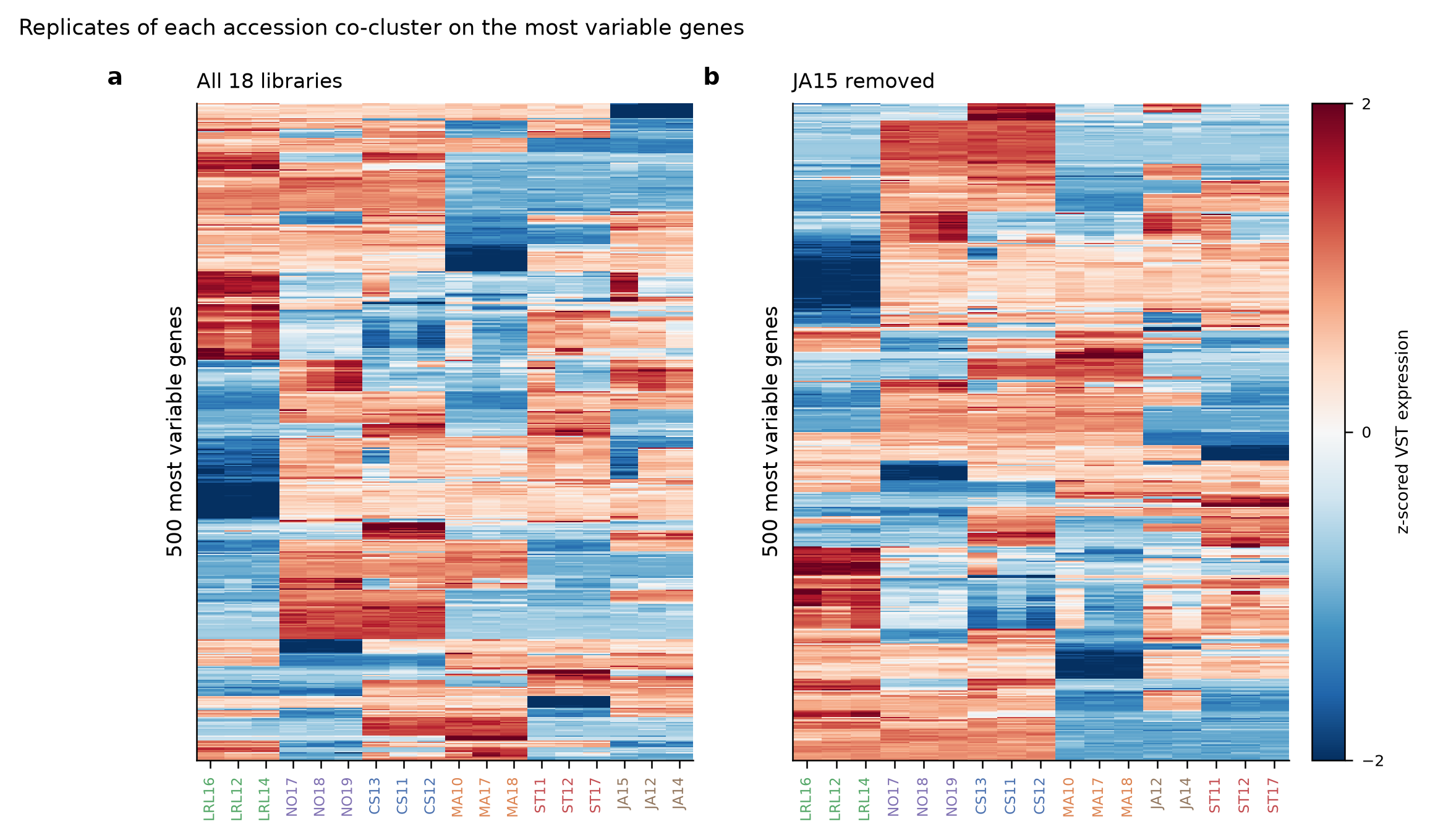

**Figure S2.** Expression heatmap of the 500 most variable genes. Z-scored variance-stabilized expression for the 500 genes with the highest variance across the 18 libraries, with libraries ordered by correlation clustering and labels coloured by cultivar. (a) All 18 libraries. (b) The same analysis with library JA15 removed. Replicates are grouped by cultivar in both, so the grouping does not depend on any single library.

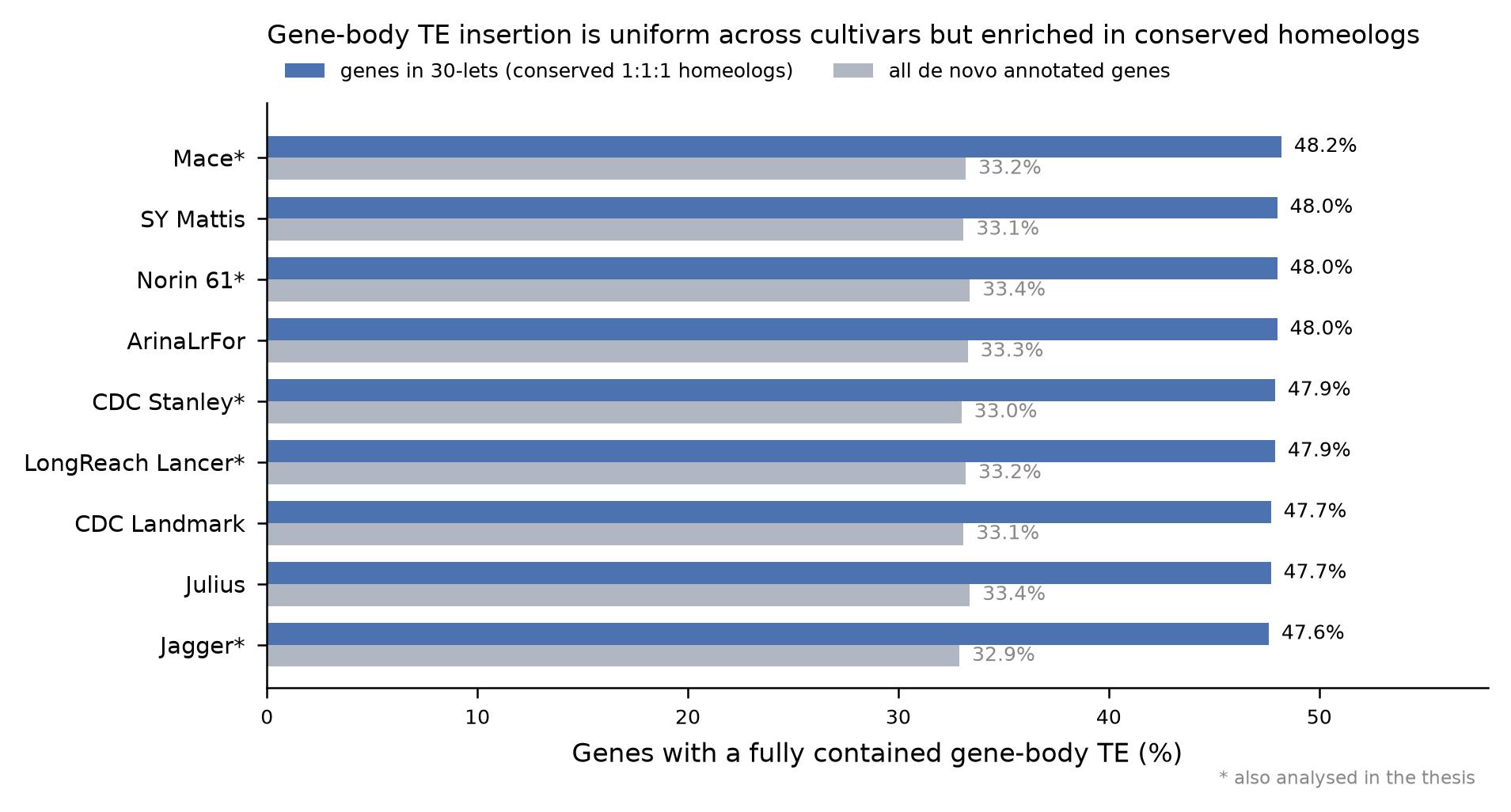

**Figure S3.** Transposable element prevalence in the wheat pan-transcriptome. Percentage of genes carrying a contained element in each of the nine sequenced cultivars, computed on each cultivar's own genome assembly, shown for all annotated genes and for the subset belonging to homoeologous groups complete in all nine cultivars. Asterisks mark the five cultivars that are also part of the six-cultivar panel analysed here.

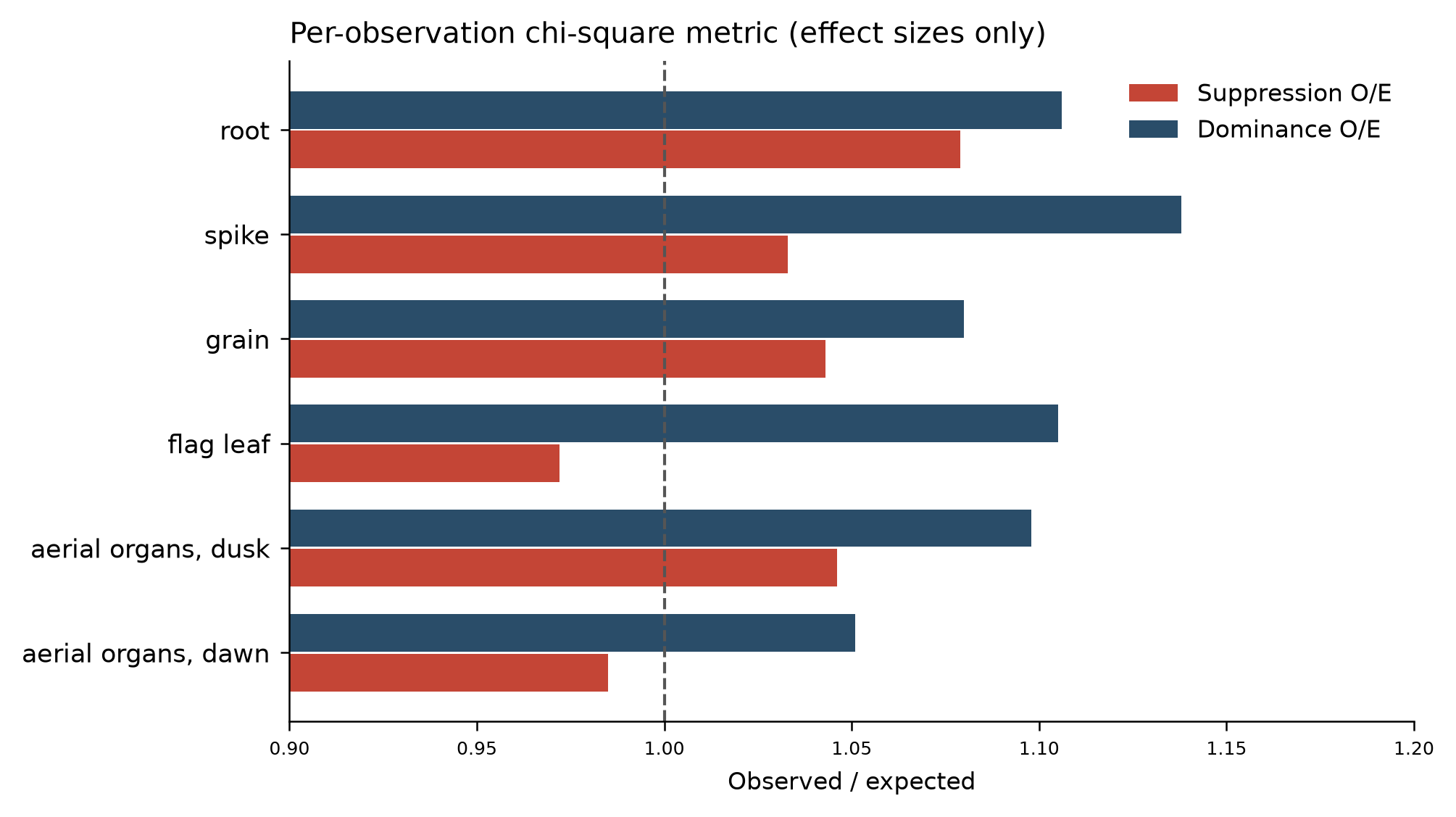

**Figure S4.** Per-tissue association in the pan-transcriptome under the per-observation chi-square metric. Observed over expected counts for the suppressed and dominant classes in each tissue. Effect sizes only are shown: because the same homoeologous group contributes an observation in every cultivar and tissue, the 80,343 asymmetric observations derive from only 5,633 distinct groups and the associated p values are anti-conservative. Cramer's V for these tests ranges from 0.045 to 0.075.

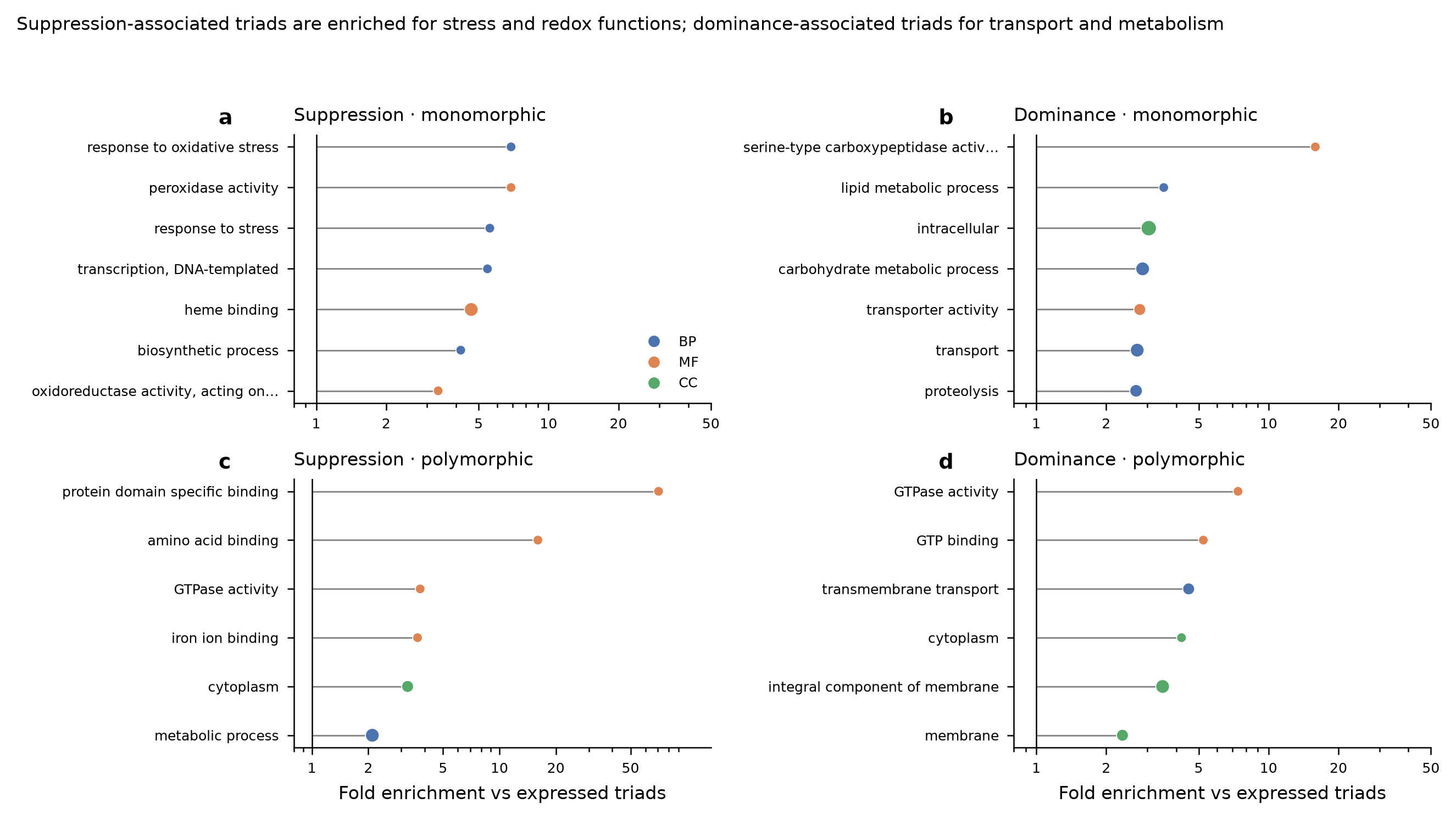

**Figure S5.** Gene Ontology enrichment of supporting groups. Terms enriched among groups supporting the suppression and dominance predictions, partitioned by whether the group was monomorphic or polymorphic for insertion and for expression. (a) Suppression, monomorphic. (b) Dominance, monomorphic. (c) Suppression, polymorphic. (d) Dominance, polymorphic. Point colour gives the Gene Ontology domain: BP, biological process; MF, molecular function; CC, cellular component. Terms are ranked by false discovery rate; no semantic reduction was applied.

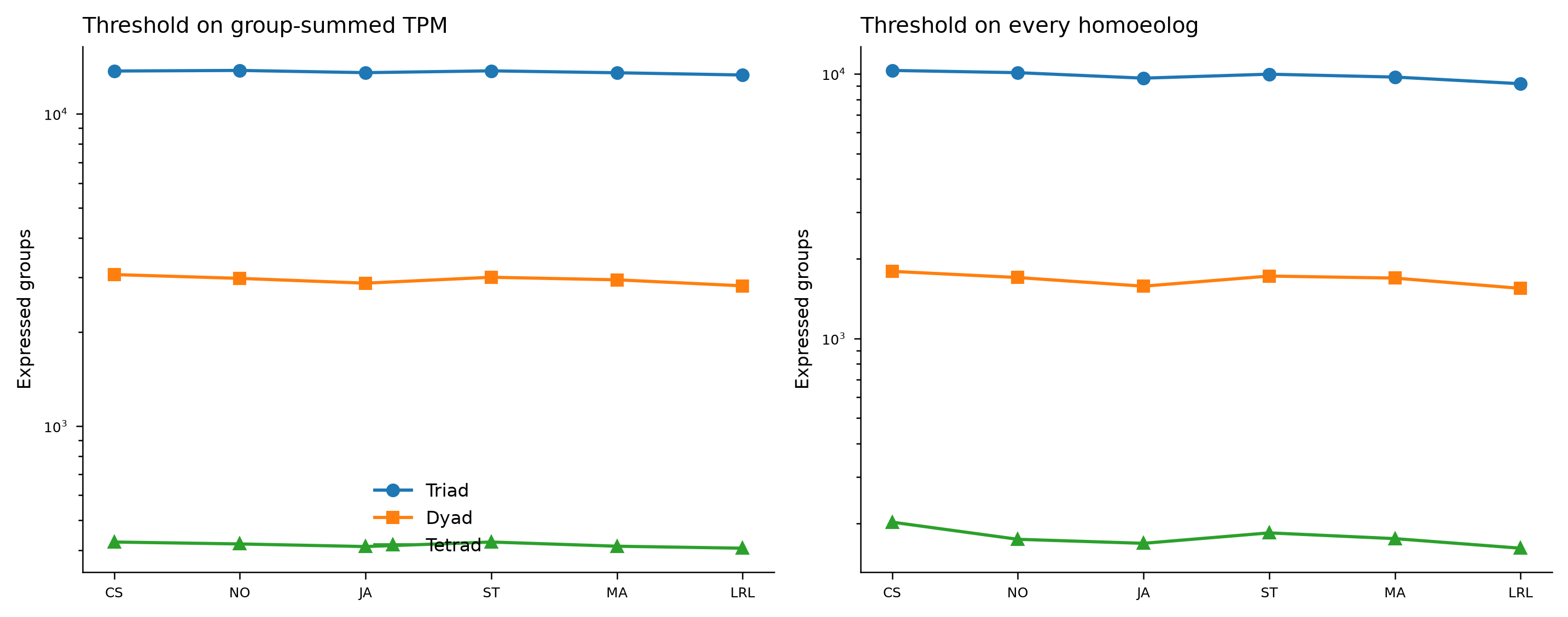

**Figure S6.** Sensitivity of expressed-group counts to the expression threshold. Number of groups scored as expressed under two alternative rules: a threshold applied to the group-summed TPM (left) and a threshold applied to every homoeolog individually (right). The stricter rule reduces counts by roughly a quarter while preserving the ranking of cultivars, so the conclusions do not depend on this choice.

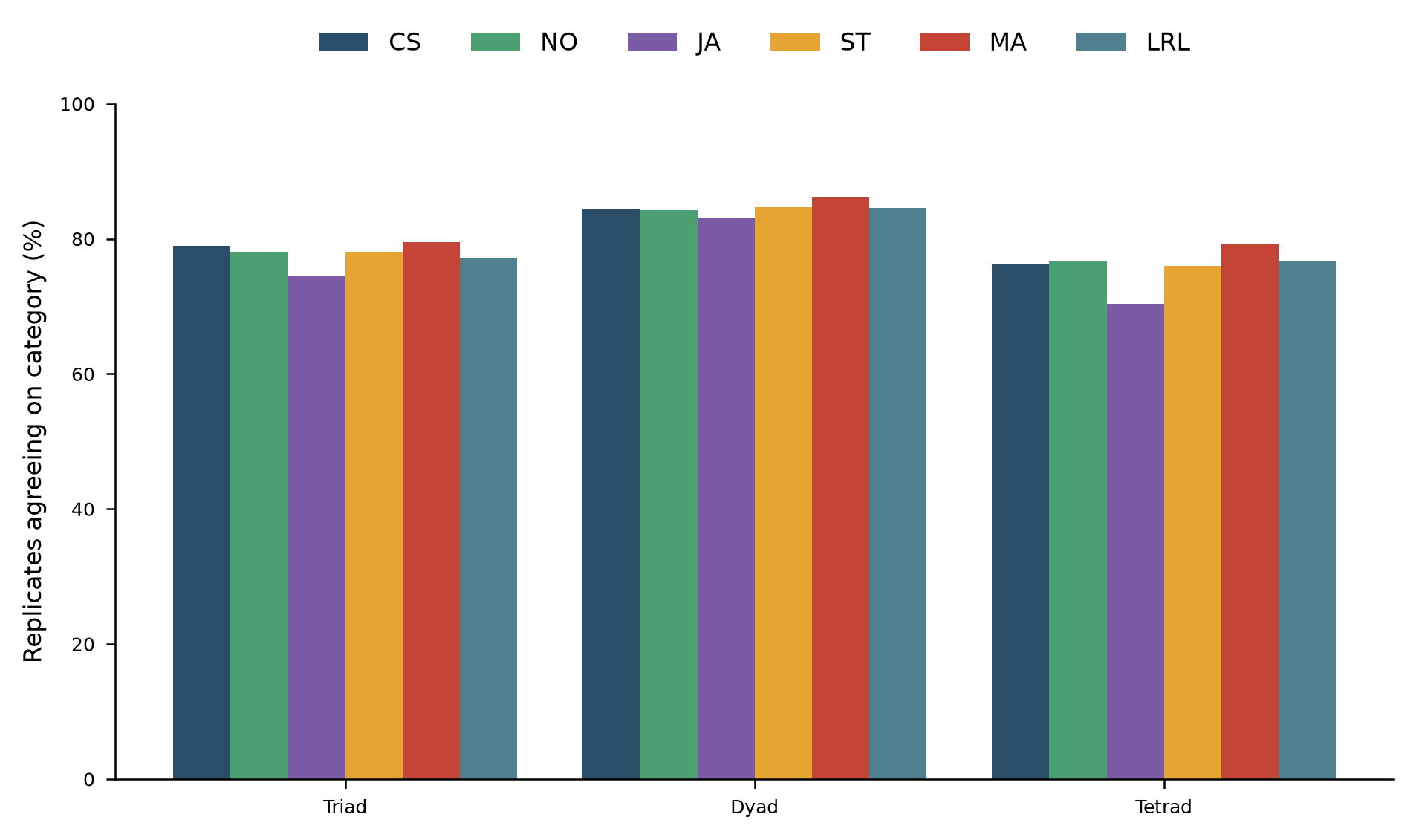

**Figure S7.** Agreement between biological replicates in expression-category assignment. For each cultivar and group type, the percentage of groups for which all three replicates were assigned the same expression category when categorised individually. Agreement is highest for dyads, which have fewer categories available, and lowest for tetrads.

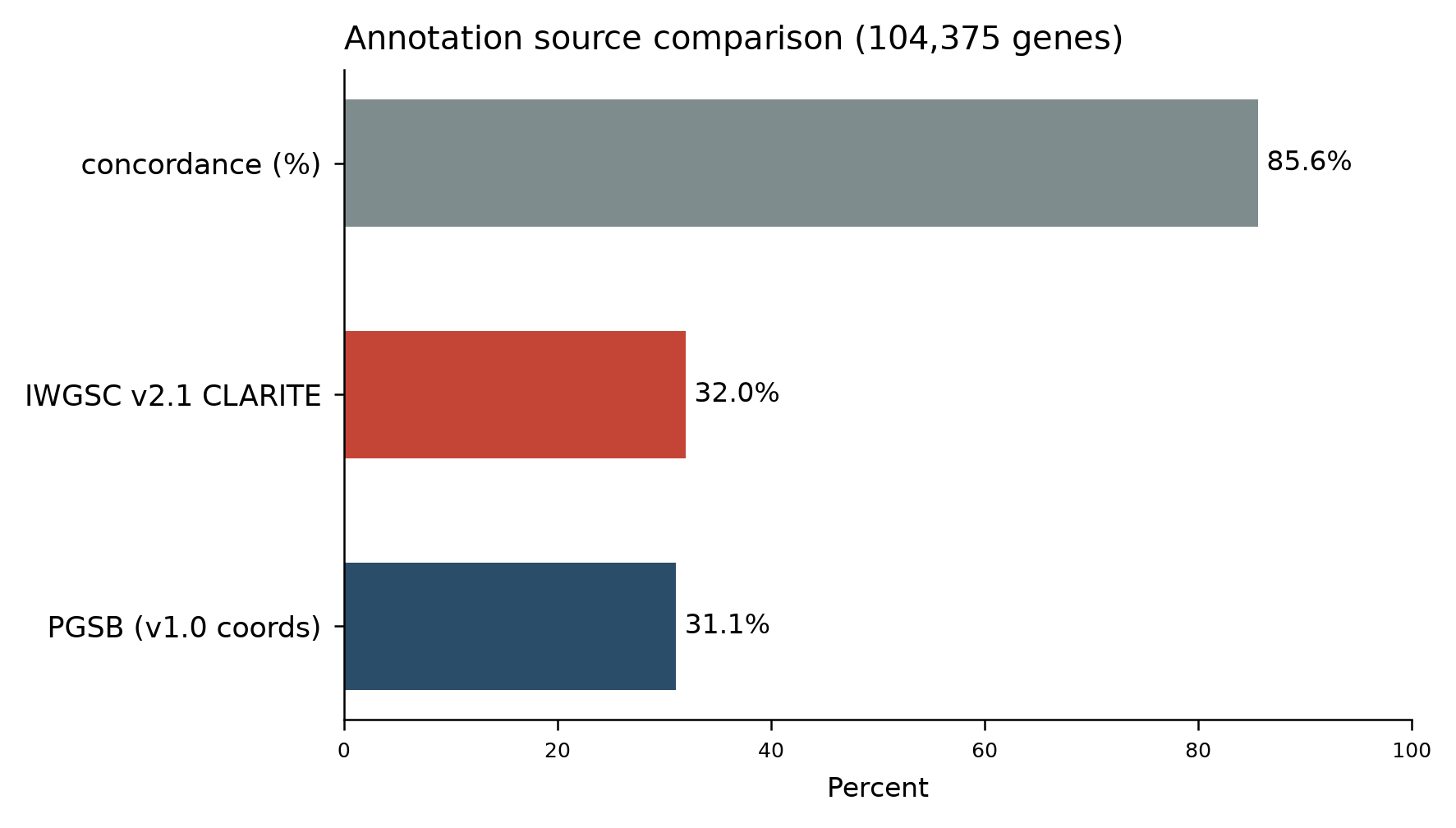

**Figure S8.** Comparison of transposable element annotation sources for Chinese Spring. Percentage of genes scored as carrying a contained element by the PGSB Triticeae-TE-library annotation used throughout this study and by the independent CLARITE annotation distributed with IWGSC RefSeq v2.1, together with the concordance between them at the level of gene-level presence or absence, across the 104,375 genes that could be compared.
