## supplemental tables for "Cultivar-specific transposable element insertions are associated with both suppression and dominance of homoeolog expression in bread wheat"

**Table S1**. Plant material, sequencing yield, and pseudoalignment. Three biological replicates were sequenced per cultivar. Read pairs are 150 bp paired-end. Pseudoalignment is against the Chinese Spring IWGSC RefSeq v2.1 transcriptome and is given as the mean ± standard deviation across the three libraries. Rates decline with genetic distance from the reference.

| **Cultivar** | **Origin** | **Libraries** | **Read pairs (M)** | **Pseudoaligned (%)** | **Uniquely assigned (%)** |
| --- | --- | --- | --- | --- | --- |
| Chinese Spring | China | 3 | 177.7 | 86.61 ± 1.24 | 59.26 ± 0.71 |
| Norin 61 | Japan | 3 | 170.6 | 84.49 ± 0.57 | 56.85 ± 0.57 |
| Jagger | USA | 3 | 180.6 | 82.20 ± 1.42 | 54.57 ± 1.21 |
| CDC Stanley | Canada | 3 | 189.5 | 81.74 ± 0.51 | 56.15 ± 0.41 |
| Mace | Australia | 3 | 196.7 | 81.50 ± 0.35 | 54.94 ± 0.87 |
| LongReach Lancer | Australia | 3 | 160.1 | 79.67 ± 0.16 | 53.79 ± 0.46 |
| Total | — | 18 | 1075.3 | — | — |

Table S2a. Homoeolog expression category counts among triads, by cultivar. Categories were assigned to each expressed triad by nearest ideal expression vector. 'Not expressed' denotes triads in which at least one homoeolog fell below the expression threshold in that cultivar.

| **Category** | **CS** | **NO** | **JA** | **ST** | **MA** | **LRL** |
| --- | --- | --- | --- | --- | --- | --- |
| Balanced | 9,304 | 9,010 | 8,575 | 8,602 | 8,418 | 7,709 |
| A dominant | 387 | 412 | 401 | 422 | 443 | 482 |
| B dominant | 467 | 433 | 399 | 430 | 465 | 373 |
| D dominant | 406 | 447 | 469 | 492 | 482 | 524 |
| A suppressed | 1,034 | 1,091 | 1,165 | 1,217 | 1,189 | 1,034 |
| B suppressed | 965 | 1,256 | 1,340 | 1,492 | 1,376 | 2,088 |
| D suppressed | 854 | 818 | 900 | 778 | 782 | 771 |
| not expressed | 4,947 | 4,897 | 5,115 | 4,931 | 5,209 | 5,383 |

Table S2b. Genes differentially expressed between each cultivar and Chinese Spring. Gene-level estimated counts for the 95,622 loci with at least 10 counts in at least three libraries were tested with PyDESeq2 v0.5.4 using cultivar as the only factor; genes with a Benjamini-Hochberg adjusted p-value below 0.05 and an absolute log2 fold change of at least 1 were called differentially expressed. Pseudoalignment rate is repeated from Table S1 because these counts scale with distance from the reference and are an upper bound on genuine expression divergence. Across all five contrasts, 38,091 genes (39.8% of those tested) were differentially expressed in at least one cultivar.

| **Cultivar** | **Pseudoaligned (%)** | **DE genes vs Chinese Spring** | **% of 95,622 tested** |
| --- | --- | --- | --- |
| Norin 61 | 84.49 | 9,123 | 9.5 |
| Mace | 81.50 | 14,131 | 14.8 |
| Jagger | 82.20 | 16,418 | 17.2 |
| CDC Stanley | 81.74 | 16,506 | 17.3 |
| LongReach Lancer | 79.67 | 20,654 | 21.6 |

Table S2c. Gene Ontology terms enriched or depleted among triads whose expression category varies between cultivars. The 17,217 genes of the 5,739 category-variable triads were tested against the 36,717 genes of all triads expressed in all six cultivars by Fisher's exact test with Benjamini-Hochberg correction; terms annotated to fewer than ten background genes were not tested. All 136 terms significant at a false discovery rate of 5% are listed, of 614 tested, ranked by q. Fold values below 1 indicate depletion. BP, biological process; MF, molecular function; CC, cellular component.

| **GO ID** | **Term** | **Domain** | **Genes in study set** | **Genes in background** | **Expected** | **Fold** | **q** |
| --- | --- | --- | --- | --- | --- | --- | --- |
| GO:0004553 | hydrolase activity, hydrolyzing O-glycosyl compounds | MF | 237 | 325 | 148.1 | 1.60 | 8.0e-21 |
| GO:0005975 | carbohydrate metabolic process | BP | 355 | 550 | 250.6 | 1.42 | 5.4e-17 |
| GO:0005215 | transporter activity | MF | 264 | 418 | 190.5 | 1.39 | 8.2e-11 |
| GO:0005488 | binding | MF | 179 | 576 | 262.4 | 0.68 | 1.2e-10 |
| GO:0016758 | transferase activity, transferring hexosyl groups | MF | 127 | 178 | 81.1 | 1.57 | 4.5e-10 |
| GO:0055114 | oxidation-reduction process | BP | 862 | 1599 | 728.5 | 1.18 | 5.4e-10 |
| GO:0016491 | oxidoreductase activity | MF | 556 | 994 | 452.9 | 1.23 | 2.0e-09 |
| GO:0005622 | intracellular | CC | 211 | 632 | 288.0 | 0.73 | 2.9e-08 |
| GO:0005840 | ribosome | CC | 161 | 482 | 219.6 | 0.73 | 3.8e-06 |
| GO:0006412 | translation | BP | 173 | 509 | 231.9 | 0.75 | 6.1e-06 |
| GO:0022857 | transmembrane transporter activity | MF | 78 | 111 | 50.6 | 1.54 | 8.7e-06 |
| GO:0003735 | structural constituent of ribosome | MF | 173 | 506 | 230.5 | 0.75 | 9.7e-06 |
| GO:0006457 | protein folding | BP | 69 | 237 | 108.0 | 0.64 | 1.2e-05 |
| GO:0015079 | potassium ion transmembrane transporter activity | MF | 45 | 57 | 26.0 | 1.73 | 1.5e-05 |
| GO:0020037 | heme binding | MF | 198 | 335 | 152.6 | 1.30 | 2.4e-05 |
| GO:0004144 | diacylglycerol O-acyltransferase activity | MF | 18 | 18 | 8.2 | 2.19 | 2.7e-05 |
| GO:0006886 | intracellular protein transport | BP | 75 | 248 | 113.0 | 0.66 | 3.3e-05 |
| GO:0022891 | substrate-specific transmembrane transporter activity | MF | 66 | 93 | 42.4 | 1.56 | 3.3e-05 |
| GO:0046983 | protein dimerization activity | MF | 137 | 221 | 100.7 | 1.36 | 3.8e-05 |
| GO:0016788 | hydrolase activity, acting on ester bonds | MF | 82 | 122 | 55.6 | 1.48 | 4.9e-05 |
| GO:0042393 | histone binding | MF | 3 | 36 | 16.4 | 0.18 | 5.7e-05 |
| GO:0051082 | unfolded protein binding | MF | 36 | 138 | 62.9 | 0.57 | 8.0e-05 |
| GO:0008654 | phospholipid biosynthetic process | BP | 21 | 23 | 10.5 | 2.00 | 1.7e-04 |
| GO:0016157 | sucrose synthase activity | MF | 15 | 15 | 6.8 | 2.19 | 1.9e-04 |
| GO:0048544 | recognition of pollen | BP | 46 | 62 | 28.2 | 1.63 | 2.0e-04 |
| GO:0003723 | RNA binding | MF | 207 | 568 | 258.8 | 0.80 | 2.1e-04 |
| GO:0071805 | potassium ion transmembrane transport | BP | 48 | 66 | 30.1 | 1.60 | 2.2e-04 |
| GO:0016702 | oxidoreductase activity, acting on single donors with incorporation of molecular oxygen, incorporation of two atoms of oxygen | MF | 33 | 42 | 19.1 | 1.72 | 3.4e-04 |
| GO:0009086 | methionine biosynthetic process | BP | 14 | 14 | 6.4 | 2.19 | 3.4e-04 |
| GO:0030247 | polysaccharide binding | MF | 41 | 55 | 25.1 | 1.64 | 3.4e-04 |
| GO:0003676 | nucleic acid binding | MF | 516 | 1296 | 590.5 | 0.87 | 3.8e-04 |
| GO:0043531 | ADP binding | MF | 92 | 146 | 66.5 | 1.38 | 5.0e-04 |
| GO:0008137 | NADH dehydrogenase (ubiquinone) activity | MF | 0 | 18 | 8.2 | 0.00 | 6.1e-04 |
| GO:0008235 | metalloexopeptidase activity | MF | 0 | 18 | 8.2 | 0.00 | 6.1e-04 |
| GO:0004499 | N,N-dimethylaniline monooxygenase activity | MF | 13 | 13 | 5.9 | 2.19 | 6.1e-04 |
| GO:0009269 | response to desiccation | BP | 13 | 13 | 5.9 | 2.19 | 6.1e-04 |
| GO:0007018 | microtubule-based movement | BP | 45 | 63 | 28.7 | 1.57 | 6.1e-04 |
| GO:0003777 | microtubule motor activity | MF | 45 | 63 | 28.7 | 1.57 | 6.1e-04 |
| GO:0005506 | iron ion binding | MF | 183 | 322 | 146.7 | 1.25 | 7.4e-04 |
| GO:0050662 | coenzyme binding | MF | 61 | 91 | 41.5 | 1.47 | 7.4e-04 |
| GO:0008483 | transaminase activity | MF | 36 | 48 | 21.9 | 1.65 | 8.5e-04 |
| GO:0008017 | microtubule binding | MF | 60 | 90 | 41.0 | 1.46 | 1.0e-03 |
| GO:0009916 | alternative oxidase activity | MF | 12 | 12 | 5.5 | 2.19 | 1.0e-03 |
| GO:0045017 | glycerolipid biosynthetic process | BP | 12 | 12 | 5.5 | 2.19 | 1.0e-03 |
| GO:0016780 | phosphotransferase activity, for other substituted phosphate groups | MF | 12 | 12 | 5.5 | 2.19 | 1.0e-03 |
| GO:0006855 | drug transmembrane transport | BP | 48 | 69 | 31.4 | 1.53 | 1.0e-03 |
| GO:0015297 | antiporter activity | MF | 48 | 69 | 31.4 | 1.53 | 1.0e-03 |
| GO:0015238 | drug transmembrane transporter activity | MF | 48 | 69 | 31.4 | 1.53 | 1.0e-03 |
| GO:0005515 | protein binding | MF | 2250 | 5212 | 2374.7 | 0.95 | 1.2e-03 |
| GO:0015935 | small ribosomal subunit | CC | 3 | 29 | 13.2 | 0.23 | 1.2e-03 |
| GO:0016021 | integral component of membrane | CC | 664 | 1307 | 595.5 | 1.12 | 1.2e-03 |
| GO:0016117 | carotenoid biosynthetic process | BP | 0 | 15 | 6.8 | 0.00 | 1.3e-03 |
| GO:0016651 | oxidoreductase activity, acting on NAD(P)H | MF | 0 | 15 | 6.8 | 0.00 | 1.3e-03 |
| GO:0016020 | membrane | CC | 717 | 1420 | 647.0 | 1.11 | 1.5e-03 |
| GO:0008641 | small protein activating enzyme activity | MF | 6 | 39 | 17.8 | 0.34 | 1.5e-03 |
| GO:0004812 | aminoacyl-tRNA ligase activity | MF | 34 | 120 | 54.7 | 0.62 | 1.6e-03 |
| GO:0006952 | defense response | BP | 24 | 30 | 13.7 | 1.76 | 1.7e-03 |
| GO:0009058 | biosynthetic process | BP | 57 | 180 | 82.0 | 0.70 | 1.7e-03 |
| GO:0036459 | thiol-dependent ubiquitinyl hydrolase activity | MF | 15 | 66 | 30.1 | 0.50 | 1.7e-03 |
| GO:0016579 | protein deubiquitination | BP | 20 | 80 | 36.5 | 0.55 | 1.8e-03 |
| GO:0016571 | histone methylation | BP | 0 | 14 | 6.4 | 0.00 | 2.2e-03 |
| GO:0004045 | aminoacyl-tRNA hydrolase activity | MF | 18 | 21 | 9.6 | 1.88 | 2.3e-03 |
| GO:0008236 | serine-type peptidase activity | MF | 53 | 80 | 36.5 | 1.45 | 2.6e-03 |
| GO:0009408 | response to heat | BP | 3 | 27 | 12.3 | 0.24 | 2.7e-03 |
| GO:0004177 | aminopeptidase activity | MF | 3 | 27 | 12.3 | 0.24 | 2.7e-03 |
| GO:0031966 | mitochondrial membrane | CC | 10 | 10 | 4.6 | 2.19 | 3.5e-03 |
| GO:0045893 | positive regulation of transcription, DNA-templated | BP | 10 | 10 | 4.6 | 2.19 | 3.5e-03 |
| GO:0015991 | ATP hydrolysis coupled proton transport | BP | 9 | 45 | 20.5 | 0.44 | 4.2e-03 |
| GO:0016779 | nucleotidyltransferase activity | MF | 12 | 54 | 24.6 | 0.49 | 4.5e-03 |
| GO:0006811 | ion transport | BP | 12 | 54 | 24.6 | 0.49 | 4.5e-03 |
| GO:0006397 | mRNA processing | BP | 14 | 60 | 27.3 | 0.51 | 5.3e-03 |
| GO:0004478 | methionine adenosyltransferase activity | MF | 0 | 12 | 5.5 | 0.00 | 5.7e-03 |
| GO:0004792 | thiosulfate sulfurtransferase activity | MF | 0 | 12 | 5.5 | 0.00 | 5.7e-03 |
| GO:0006556 | S-adenosylmethionine biosynthetic process | BP | 0 | 12 | 5.5 | 0.00 | 5.7e-03 |
| GO:0006914 | autophagy | BP | 0 | 12 | 5.5 | 0.00 | 5.7e-03 |
| GO:0004407 | histone deacetylase activity | MF | 0 | 12 | 5.5 | 0.00 | 5.7e-03 |
| GO:0004061 | arylformamidase activity | MF | 0 | 12 | 5.5 | 0.00 | 5.7e-03 |
| GO:0016575 | histone deacetylation | BP | 0 | 12 | 5.5 | 0.00 | 5.7e-03 |
| GO:0046961 | proton-transporting ATPase activity, rotational mechanism | MF | 0 | 12 | 5.5 | 0.00 | 5.7e-03 |
| GO:0019441 | tryptophan catabolic process to kynurenine | BP | 0 | 12 | 5.5 | 0.00 | 5.7e-03 |
| GO:0003918 | DNA topoisomerase type II (ATP-hydrolyzing) activity | MF | 0 | 12 | 5.5 | 0.00 | 5.7e-03 |
| GO:0019843 | rRNA binding | MF | 3 | 24 | 10.9 | 0.27 | 5.7e-03 |
| GO:0005741 | mitochondrial outer membrane | CC | 3 | 24 | 10.9 | 0.27 | 5.7e-03 |
| GO:0033014 | tetrapyrrole biosynthetic process | BP | 3 | 24 | 10.9 | 0.27 | 5.7e-03 |
| GO:0048500 | signal recognition particle | CC | 3 | 24 | 10.9 | 0.27 | 5.7e-03 |
| GO:0030131 | clathrin adaptor complex | CC | 3 | 24 | 10.9 | 0.27 | 5.7e-03 |
| GO:0006396 | RNA processing | BP | 39 | 127 | 57.9 | 0.67 | 6.1e-03 |
| GO:0016705 | oxidoreductase activity, acting on paired donors, with incorporation or reduction of molecular oxygen | MF | 135 | 240 | 109.4 | 1.23 | 6.1e-03 |
| GO:0004601 | peroxidase activity | MF | 60 | 96 | 43.7 | 1.37 | 6.5e-03 |
| GO:0005525 | GTP binding | MF | 153 | 407 | 185.4 | 0.83 | 7.5e-03 |
| GO:0015031 | protein transport | BP | 21 | 77 | 35.1 | 0.60 | 8.2e-03 |
| GO:0004519 | endonuclease activity | MF | 33 | 48 | 21.9 | 1.51 | 8.6e-03 |
| GO:0030170 | pyridoxal phosphate binding | MF | 81 | 136 | 62.0 | 1.31 | 8.6e-03 |
| GO:0006979 | response to oxidative stress | BP | 63 | 102 | 46.5 | 1.36 | 8.6e-03 |
| GO:0030117 | membrane coat | CC | 6 | 33 | 15.0 | 0.40 | 8.8e-03 |
| GO:0015986 | ATP synthesis coupled proton transport | BP | 6 | 33 | 15.0 | 0.40 | 8.8e-03 |
| GO:0005985 | sucrose metabolic process | BP | 15 | 18 | 8.2 | 1.83 | 9.1e-03 |
| GO:0033897 | ribonuclease T2 activity | MF | 15 | 18 | 8.2 | 1.83 | 9.1e-03 |
| GO:0004190 | aspartic-type endopeptidase activity | MF | 45 | 69 | 31.4 | 1.43 | 9.1e-03 |
| GO:0004674 | protein serine/threonine kinase activity | MF | 63 | 103 | 46.9 | 1.34 | 1.2e-02 |
| GO:0000166 | nucleotide binding | MF | 90 | 251 | 114.4 | 0.79 | 1.3e-02 |
| GO:0008536 | Ran GTPase binding | MF | 9 | 41 | 18.7 | 0.48 | 1.5e-02 |
| GO:0016614 | oxidoreductase activity, acting on CH-OH group of donors | MF | 39 | 60 | 27.3 | 1.43 | 1.6e-02 |
| GO:0016791 | phosphatase activity | MF | 27 | 90 | 41.0 | 0.66 | 1.7e-02 |
| GO:0003690 | double-stranded DNA binding | MF | 63 | 105 | 47.8 | 1.32 | 1.8e-02 |
| GO:0006418 | tRNA aminoacylation for protein translation | BP | 34 | 108 | 49.2 | 0.69 | 2.0e-02 |
| GO:0006265 | DNA topological change | BP | 3 | 21 | 9.6 | 0.31 | 2.0e-02 |
| GO:0046933 | proton-transporting ATP synthase activity, rotational mechanism | MF | 3 | 21 | 9.6 | 0.31 | 2.0e-02 |
| GO:0004003 | ATP-dependent DNA helicase activity | MF | 3 | 21 | 9.6 | 0.31 | 2.0e-02 |
| GO:0006367 | transcription initiation from RNA polymerase II promoter | BP | 3 | 21 | 9.6 | 0.31 | 2.0e-02 |
| GO:0033179 | proton-transporting V-type ATPase, V0 domain | CC | 3 | 21 | 9.6 | 0.31 | 2.0e-02 |
| GO:0032040 | small-subunit processome | CC | 3 | 21 | 9.6 | 0.31 | 2.0e-02 |
| GO:0005694 | chromosome | CC | 6 | 30 | 13.7 | 0.44 | 2.8e-02 |
| GO:0006888 | ER to Golgi vesicle-mediated transport | BP | 6 | 30 | 13.7 | 0.44 | 2.8e-02 |
| GO:0055085 | transmembrane transport | BP | 390 | 772 | 351.7 | 1.11 | 2.8e-02 |
| GO:0003899 | DNA-directed 5'-3' RNA polymerase activity | MF | 30 | 96 | 43.7 | 0.69 | 2.8e-02 |
| GO:0009507 | chloroplast | CC | 9 | 39 | 17.8 | 0.51 | 2.9e-02 |
| GO:0043039 | tRNA aminoacylation | BP | 9 | 39 | 17.8 | 0.51 | 2.9e-02 |
| GO:0051536 | iron-sulfur cluster binding | MF | 42 | 126 | 57.4 | 0.73 | 3.6e-02 |
| GO:0004672 | protein kinase activity | MF | 750 | 1535 | 699.4 | 1.07 | 3.8e-02 |
| GO:0006351 | transcription, DNA-templated | BP | 51 | 147 | 67.0 | 0.76 | 4.0e-02 |
| GO:0005737 | cytoplasm | CC | 212 | 532 | 242.4 | 0.87 | 4.0e-02 |
| GO:0005634 | nucleus | CC | 330 | 805 | 366.8 | 0.90 | 4.0e-02 |
| GO:0045735 | nutrient reservoir activity | MF | 12 | 15 | 6.8 | 1.76 | 4.0e-02 |
| GO:0045492 | xylan biosynthetic process | BP | 12 | 15 | 6.8 | 1.76 | 4.0e-02 |
| GO:0005759 | mitochondrial matrix | CC | 12 | 15 | 6.8 | 1.76 | 4.0e-02 |
| GO:0003978 | UDP-glucose 4-epimerase activity | MF | 12 | 15 | 6.8 | 1.76 | 4.0e-02 |
| GO:0070008 | serine-type exopeptidase activity | MF | 12 | 15 | 6.8 | 1.76 | 4.0e-02 |
| GO:0046373 | L-arabinose metabolic process | BP | 12 | 15 | 6.8 | 1.76 | 4.0e-02 |
| GO:0046556 | alpha-L-arabinofuranosidase activity | MF | 12 | 15 | 6.8 | 1.76 | 4.0e-02 |
| GO:0030151 | molybdenum ion binding | MF | 12 | 15 | 6.8 | 1.76 | 4.0e-02 |
| GO:0004707 | MAP kinase activity | MF | 8 | 34 | 15.5 | 0.52 | 4.5e-02 |
| GO:0008289 | lipid binding | MF | 45 | 74 | 33.7 | 1.33 | 4.5e-02 |
| GO:0009072 | aromatic amino acid family metabolic process | BP | 3 | 19 | 8.7 | 0.35 | 4.6e-02 |
| GO:0006468 | protein phosphorylation | BP | 750 | 1539 | 701.2 | 1.07 | 4.7e-02 |
| GO:0051539 | 4 iron, 4 sulfur cluster binding | MF | 12 | 45 | 20.5 | 0.59 | 4.9e-02 |

Table S3. Transposable element insertion state by homoeologous group type. A group was scored as carrying an element when at least one cultivar carried an element entirely within the gene body of at least one member gene.

| **group_type** | **with TE in >=1 cultivar** | **without TE in any cultivar** | **% with TE** |
| --- | --- | --- | --- |
| Dyad | 1898 | 2683 | 41.4 |
| Tetrad | 202 | 255 | 44.2 |
| Triad | 9435 | 6650 | 58.7 |

Table S4. Position of contained transposable elements by superfamily, pooled across the six cultivars. An element was classed as overlapping coding sequence when any part of it intersected an annotated coding exon.

| **Superfamily** | **Code** | **Contained elements** | **% overlapping CDS** |
| --- | --- | --- | --- |
| Gypsy | RLG | 94,943 | 23.01 |
| CACTA | DTC | 80,210 | 22.10 |
| Copia | RLC | 79,560 | 18.98 |
| LTR, unclassified | RLX | 169,647 | 17.53 |
| Unknown | XXX | 29,718 | 7.35 |
| Mutator | DTM | 8,095 | 6.89 |
| Harbinger | DTH | 6,073 | 3.56 |
| LINE | RIX | 23,900 | 3.50 |
| Unclassified | DXX | 74,383 | 3.08 |
| Tc1/Mariner | DTT | 28,100 | 1.92 |
| DNA, unclassified | DTX | 14,880 | 1.61 |

Table S5. Per-superfamily association statistics for all group types and both predictions. Insertion patterns were rebuilt from elements of a single superfamily at a time and tested against the same permutation null. 'All elements' gives the unrestricted test for comparison. Obs., observed supporting groups; Exp., mean expected across 2,000 permutations; p, empirical p value floored at 1/2001; q, Benjamini-Hochberg corrected.

| **Superfamily** | **Genes** | **Group type** | **Prediction** | **Obs.** | **Exp.** | **Fold** | **p** | **q** |
| --- | --- | --- | --- | --- | --- | --- | --- | --- |
| All elements | 39,872 | Triad | suppression | 183 | 115.4 | 1.586 | 0.001 | 0.005 |
| All elements | 39,872 | Triad | dominance | 171 | 154.3 | 1.108 | 0.154 | 0.246 |
| All elements | 39,872 | Dyad | suppression | 37 | 25.5 | 1.452 | 0.023 | 0.120 |
| All elements | 39,872 | Dyad | dominance | 28 | 25.1 | 1.114 | 0.577 | 0.865 |
| All elements | 39,872 | Tetrad | suppression | 5 | 5.7 | 0.880 | 0.976 | 1.000 |
| All elements | 39,872 | Tetrad | dominance | 4 | 5.3 | 0.756 | 0.746 | 1.000 |
| LTR, unclass. | 13,264 | Triad | suppression | 138 | 105.7 | 1.306 | 0.001 | 0.005 |
| LTR, unclass. | 13,264 | Triad | dominance | 124 | 134.0 | 0.925 | 0.393 | 0.432 |
| LTR, unclass. | 13,264 | Dyad | suppression | 27 | 18.4 | 1.470 | 0.054 | 0.216 |
| LTR, unclass. | 13,264 | Dyad | dominance | 19 | 17.8 | 1.067 | 0.843 | 1.000 |
| LTR, unclass. | 13,264 | Tetrad | suppression | 5 | 5.0 | 1.002 | 1.000 | 1.000 |
| LTR, unclass. | 13,264 | Tetrad | dominance | 4 | 4.1 | 0.983 | 1.000 | 1.000 |
| Unclass. | 9,221 | Triad | suppression | 107 | 78.2 | 1.368 | 0.001 | 0.005 |
| Unclass. | 9,221 | Triad | dominance | 97 | 101.9 | 0.952 | 0.630 | 0.657 |
| Unclass. | 9,221 | Dyad | suppression | 14 | 13.5 | 1.040 | 0.946 | 1.000 |
| Unclass. | 9,221 | Dyad | dominance | 14 | 13.5 | 1.038 | 0.974 | 1.000 |
| Unclass. | 9,221 | Tetrad | suppression | 3 | 2.8 | 1.059 | 1.000 | 1.000 |
| Unclass. | 9,221 | Tetrad | dominance | 5 | 2.4 | 2.073 | 0.149 | 1.000 |
| Gypsy | 7,811 | Triad | suppression | 94 | 70.7 | 1.330 | 0.008 | 0.027 |
| Gypsy | 7,811 | Triad | dominance | 72 | 89.6 | 0.804 | 0.054 | 0.118 |
| Gypsy | 7,811 | Dyad | suppression | 23 | 14.1 | 1.632 | 0.025 | 0.120 |
| Gypsy | 7,811 | Dyad | dominance | 14 | 13.4 | 1.047 | 0.928 | 1.000 |
| Gypsy | 7,811 | Tetrad | suppression | 1 | 3.5 | 0.289 | 0.234 | 1.000 |
| Gypsy | 7,811 | Tetrad | dominance | 6 | 3.1 | 1.932 | 0.150 | 1.000 |
| CACTA | 7,779 | Triad | suppression | 79 | 71.4 | 1.106 | 0.396 | 0.432 |
| CACTA | 7,779 | Triad | dominance | 73 | 90.8 | 0.804 | 0.047 | 0.113 |
| CACTA | 7,779 | Dyad | suppression | 18 | 11.6 | 1.553 | 0.076 | 0.249 |
| CACTA | 7,779 | Dyad | dominance | 9 | 11.8 | 0.762 | 0.491 | 0.841 |
| CACTA | 7,779 | Tetrad | suppression | 4 | 4.8 | 0.841 | 0.967 | 1.000 |
| CACTA | 7,779 | Tetrad | dominance | 5 | 4.4 | 1.148 | 0.875 | 1.000 |
| Copia | 6,325 | Triad | suppression | 95 | 59.2 | 1.605 | 0.001 | 0.005 |
| Copia | 6,325 | Triad | dominance | 54 | 73.7 | 0.733 | 0.011 | 0.033 |
| Copia | 6,325 | Dyad | suppression | 22 | 9.5 | 2.317 | 0.001 | 0.024 |
| Copia | 6,325 | Dyad | dominance | 9 | 9.7 | 0.927 | 0.982 | 1.000 |
| Copia | 6,325 | Tetrad | suppression | 2 | 3.2 | 0.628 | 0.698 | 1.000 |
| Copia | 6,325 | Tetrad | dominance | 4 | 3.0 | 1.347 | 0.695 | 1.000 |
| Tc1/Mariner | 4,309 | Triad | suppression | 53 | 42.2 | 1.255 | 0.105 | 0.210 |
| Tc1/Mariner | 4,309 | Triad | dominance | 46 | 53.8 | 0.856 | 0.313 | 0.395 |
| Tc1/Mariner | 4,309 | Dyad | suppression | 10 | 6.9 | 1.451 | 0.298 | 0.596 |
| Tc1/Mariner | 4,309 | Dyad | dominance | 10 | 7.5 | 1.338 | 0.395 | 0.729 |
| Tc1/Mariner | 4,309 | Tetrad | suppression | 1 | 1.2 | 0.861 | 1.000 | 1.000 |
| Tc1/Mariner | 4,309 | Tetrad | dominance | 0 | 1.0 | 0.000 | 0.674 | 1.000 |
| Unknown | 3,038 | Triad | suppression | 49 | 31.5 | 1.555 | 0.001 | 0.005 |
| Unknown | 3,038 | Triad | dominance | 33 | 38.9 | 0.848 | 0.363 | 0.432 |
| Unknown | 3,038 | Dyad | suppression | 11 | 6.2 | 1.777 | 0.095 | 0.253 |
| Unknown | 3,038 | Dyad | dominance | 2 | 6.3 | 0.316 | 0.083 | 0.249 |
| Unknown | 3,038 | Tetrad | suppression | 0 | 0.8 | 0.000 | 0.825 | 1.000 |
| Unknown | 3,038 | Tetrad | dominance | 1 | 0.7 | 1.464 | 1.000 | 1.000 |
| LINE | 2,329 | Triad | suppression | 30 | 24.7 | 1.213 | 0.313 | 0.395 |
| LINE | 2,329 | Triad | dominance | 23 | 31.2 | 0.737 | 0.152 | 0.246 |
| LINE | 2,329 | Dyad | suppression | 12 | 4.3 | 2.764 | 0.005 | 0.048 |
| LINE | 2,329 | Dyad | dominance | 1 | 4.3 | 0.233 | 0.111 | 0.266 |
| LINE | 2,329 | Tetrad | suppression | 0 | 1.8 | 0.000 | 0.269 | 1.000 |
| LINE | 2,329 | Tetrad | dominance | 0 | 1.6 | 0.000 | 0.364 | 1.000 |
| DNA, unclass. | 2,170 | Triad | suppression | 32 | 24.4 | 1.309 | 0.133 | 0.245 |
| DNA, unclass. | 2,170 | Triad | dominance | 24 | 30.6 | 0.784 | 0.249 | 0.351 |
| DNA, unclass. | 2,170 | Dyad | suppression | 7 | 3.5 | 1.973 | 0.123 | 0.268 |
| DNA, unclass. | 2,170 | Dyad | dominance | 5 | 3.5 | 1.417 | 0.561 | 0.865 |
| DNA, unclass. | 2,170 | Tetrad | suppression | 1 | 0.5 | 1.974 | 0.864 | 1.000 |
| DNA, unclass. | 2,170 | Tetrad | dominance | 0 | 0.4 | 0.000 | 1.000 | 1.000 |
| Mutator | 871 | Triad | suppression | 15 | 6.4 | 2.359 | 0.005 | 0.020 |
| Mutator | 871 | Triad | dominance | 12 | 8.1 | 1.486 | 0.224 | 0.336 |
| Mutator | 871 | Dyad | suppression | 7 | 1.8 | 3.960 | 0.006 | 0.048 |
| Mutator | 871 | Dyad | dominance | 1 | 1.9 | 0.518 | 0.818 | 1.000 |
| Mutator | 871 | Tetrad | suppression | 0 | 0.3 | 0.000 | 1.000 | 1.000 |
| Mutator | 871 | Tetrad | dominance | 0 | 0.4 | 0.000 | 1.000 | 1.000 |
| Harbinger | 687 | Triad | suppression | 6 | 7.5 | 0.795 | 0.722 | 0.722 |
| Harbinger | 687 | Triad | dominance | 17 | 9.4 | 1.805 | 0.020 | 0.053 |
| Harbinger | 687 | Dyad | suppression | 1 | 1.5 | 0.665 | 1.000 | 1.000 |
| Harbinger | 687 | Dyad | dominance | 2 | 1.6 | 1.287 | 0.954 | 1.000 |
| Harbinger | 687 | Tetrad | suppression | 1 | 0.7 | 1.439 | 1.000 | 1.000 |
| Harbinger | 687 | Tetrad | dominance | 1 | 0.5 | 1.820 | 0.901 | 1.000 |

Table S6. Pan-transcriptome association statistics under the two alignment strategies. The same reads were quantified either against the Chinese Spring reference or against each cultivar's own genome assembly, and the identical support rule and permutation null applied to each. Aligning to a single reference inflates the suppression enrichment in every tissue while leaving dominance essentially unchanged.

| **Alignment** | **Tissue** | **Prediction** | **Observed** | **Expected** | **Fold** | **q** |
| --- | --- | --- | --- | --- | --- | --- |
| Chinese Spring | Aerial organs, dawn | suppression | 97 | 66.5 | 1.458 | 0.0120 |
| Chinese Spring | Aerial organs, dawn | dominance | 92 | 81.3 | 1.131 | 0.3502 |
| Chinese Spring | Aerial organs, dusk | suppression | 86 | 56.3 | 1.527 | 0.0048 |
| Chinese Spring | Aerial organs, dusk | dominance | 84 | 70.9 | 1.185 | 0.2507 |
| Chinese Spring | Root | suppression | 99 | 60.6 | 1.633 | 0.0048 |
| Chinese Spring | Root | dominance | 95 | 81.8 | 1.162 | 0.2952 |
| Chinese Spring | Spike | suppression | 77 | 48.3 | 1.594 | 0.0048 |
| Chinese Spring | Spike | dominance | 66 | 67.6 | 0.977 | 0.9522 |
| Chinese Spring | Flag leaf | suppression | 91 | 76.7 | 1.187 | 0.2422 |
| Chinese Spring | Flag leaf | dominance | 96 | 85.6 | 1.122 | 0.3794 |
| Chinese Spring | Grain | suppression | 100 | 60.8 | 1.644 | 0.0048 |
| Chinese Spring | Grain | dominance | 78 | 80.0 | 0.975 | 0.9522 |
| own-genome | Aerial organs, dawn | suppression | 75 | 61.7 | 1.215 | 0.2422 |
| own-genome | Aerial organs, dawn | dominance | 81 | 71.9 | 1.126 | 0.3994 |
| own-genome | Aerial organs, dusk | suppression | 62 | 50.0 | 1.240 | 0.2422 |
| own-genome | Aerial organs, dusk | dominance | 70 | 60.5 | 1.157 | 0.3502 |
| own-genome | Root | suppression | 71 | 54.2 | 1.309 | 0.1097 |
| own-genome | Root | dominance | 78 | 69.5 | 1.122 | 0.4304 |
| own-genome | Spike | suppression | 52 | 42.3 | 1.230 | 0.2952 |
| own-genome | Spike | dominance | 56 | 57.7 | 0.971 | 0.9522 |
| own-genome | Flag leaf | suppression | 72 | 70.2 | 1.026 | 0.9522 |
| own-genome | Flag leaf | dominance | 81 | 73.7 | 1.099 | 0.5000 |
| own-genome | Grain | suppression | 79 | 55.7 | 1.417 | 0.0048 |
| own-genome | Grain | dominance | 71 | 70.8 | 1.002 | 1.0000 |

Table S7. Percentage of homoeologous groups assigned the identical bias class under Chinese Spring alignment and under own-genome alignment, by cultivar and tissue. Overall agreement is 90.2%.

| **Cultivar** | **Aerial organs, dawn** | **Aerial organs, dusk** | **Root** | **Spike** | **Flag leaf** | **Grain** |
| --- | --- | --- | --- | --- | --- | --- |
| ARI | 91.0 | 91.2 | 91.9 | 91.5 | 90.5 | 90.5 |
| JAG | 90.4 | 90.9 | 91.8 | 90.8 | 90.6 | 90.3 |
| JUL | 91.2 | 91.8 | 91.8 | 91.7 | 90.9 | 90.6 |
| LAC | 86.3 | 86.3 | 86.8 | 87.1 | 86.8 | 84.7 |
| LDM | 92.4 | 92.9 | 92.6 | 93.1 | 91.8 | 91.8 |
| MAC | 85.6 | 88.7 | 92.6 | 92.0 | 91.7 | nan |
| NOR | 87.5 | 88.1 | 88.3 | 87.8 | 87.3 | 87.4 |
| STA | 90.9 | 91.4 | 92.1 | 91.8 | 89.9 | nan |
| SYM | 90.5 | 90.8 | 91.4 | 91.4 | 90.1 | 88.7 |

Table S8. Complete list of candidate loci at which a polymorphic transposable element insertion coincides with suppression of the inserted homoeolog in every cultivar that carries it (73 groups, 74 candidate homoeologs). Each row names the homoeolog whose insertion status varies between cultivars. Cultivars: CS, Chinese Spring; MA, Mace; LRL, LongReach Lancer; ST, CDC Stanley; NO, Norin 61; JA, Jagger. Approximately 15% of this list is expected by chance.

| **No.** | **Gene (v2.1)** | **Chr** | **Sub** | **Annotation** | **TE present in** | **TE absent in** |
| --- | --- | --- | --- | --- | --- | --- |
| 1 | TraesCS2B03G1134600 | 2B | B | RNA binding protein | LRL | CS, MA, ST, NO, JA |
| 2 | TraesCS3D03G1115100 | 3D | D | — | LRL | CS, MA, ST, NO, JA |
| 3 | TraesCS2D03G1107100 | 2D | D | Ribosomal protein S4 | JA | CS, MA, LRL, ST, NO |
| 4 | TraesCS6D03G0856400 | 6D | D | Phosphoglycolate phosphatase, putative | CS, MA, ST, NO, JA | LRL |
| 5 | TraesCS4B03G0094800 | 4B | B | — | ST | CS, MA, LRL, NO, JA |
| 6 | TraesCS7A03G0109800 | 7A | A | — | JA | CS, MA, LRL, ST, NO |
| 7 | TraesCS1B03G0004000 | 1B | B | — | MA, LRL, ST, NO, JA | CS |
| 8 | TraesCS6B03G0359800 | 6B | B | Calcium-dependent lipid-binding domain p | ST, NO | CS, MA, LRL, JA |
| 9 | TraesCS2B03G1146400 | 2B | B | Sodium transporter | LRL | CS, MA, ST, NO, JA |
| 10 | TraesCS6A03G0043500 | 6A | A | Mitochondrial transcription termination | ST | CS, MA, LRL, NO, JA |
| 11 | TraesCS2A03G0101800 | 2A | A | Cytochrome P450 | MA, ST, JA | CS, LRL, NO |
| 12 | TraesCS5A03G0572200 | 5A | A | Zinc finger protein | CS, LRL, ST, NO, JA | MA |
| 13 | TraesCS3B03G1468900 | 3B | B | — | LRL | CS, MA, ST, NO, JA |
| 14 | TraesCS2D03G1268700 | 2D | D | Isoaspartyl peptidase/L-asparaginase | LRL | CS, MA, ST, NO, JA |
| 15 | TraesCS2B03G1456000 | 2B | B | Pentatricopeptide repeat-containing prot | ST, JA | CS, MA, LRL, NO |
| 16 | TraesCS7B03G0807500 | 7B | B | F-box/kelch-repeat protein | LRL, ST | CS, MA, NO, JA |
| 17 | TraesCS2B03G0899600 | 2B | B | ATP-dependent zinc metalloprotease FtsH | LRL | CS, MA, ST, NO, JA |
| 18 | TraesCS5A03G0672700 | 5A | A | — | LRL, ST | CS, MA, NO, JA |
| 19 | TraesCS1B03G1251400 | 1B | B | — | MA, NO | CS, LRL, ST, JA |
| 20 | TraesCS1D03G1016800 | 1D | D | RNA binding protein | JA | CS, MA, LRL, ST, NO |
| 21 | TraesCS2B03G0487500 | 2B | B | — | LRL | CS, MA, ST, NO, JA |
| 22 | TraesCS6B03G0227400 | 6B | B | Pentatricopeptide repeat-containing prot | MA | CS, LRL, ST, NO, JA |
| 23 | TraesCS5B03G1026600 | 5B | B | Hexosyltransferase | MA | CS, LRL, ST, NO, JA |
| 24 | TraesCS3D03G1154800 | 3D | D | — | LRL | CS, MA, ST, NO, JA |
| 25 | TraesCS3B03G1067900 | 3B | B | — | NO | CS, MA, LRL, ST, JA |
| 26 | TraesCS1B03G0950200 | 1B | B | Cleavage stimulation factor subunit | NO | CS, MA, LRL, ST, JA |
| 27 | TraesCS5A03G0618300 | 5A | A | Phosphoribosylaminoimidazole-succinocarb | LRL, ST, JA | CS, MA, NO |
| 28 | TraesCS5B03G1202900 | 5B | B | Ribosomal RNA large subunit methyltransf | LRL | CS, MA, ST, NO, JA |
| 29 | TraesCS2B03G1081400 | 2B | B | — | LRL | CS, MA, ST, NO, JA |
| 30 | TraesCS2B03G1374000 | 2B | B | — | LRL | CS, MA, ST, NO, JA |
| 31 | TraesCS6B03G1026600 | 6B | B | — | ST | CS, MA, LRL, NO, JA |
| 32 | TraesCS2D03G1292700 | 2D | D | Protein kinase, putative | LRL | CS, MA, ST, NO, JA |
| 33 | TraesCS2D03G1091300 | 2D | D | Cell division protein ftsZ | JA | CS, MA, LRL, ST, NO |
| 34 | TraesCS3B03G0522200 | 3B | B | Cysteine proteinase inhibitor | MA, LRL, ST, JA | CS, NO |
| 35 | TraesCS1B03G1078600 | 1B | B | tRNA-2-methylthio-N(6)-dimethylallyladen | LRL, ST, JA | CS, MA, NO |
| 36 | TraesCS2B03G0865500 | 2B | B | — | LRL | CS, MA, ST, NO, JA |
| 37 | TraesCS7B03G0157100 | 7B | B | zinc ion binding protein | MA, LRL, ST, JA | CS, NO |
| 38 | TraesCS2B03G0541200 | 2B | B | — | LRL | CS, MA, ST, NO, JA |
| 39 | TraesCS6B03G1091800 | 6B | B | Serine incorporator | LRL, ST, JA | CS, MA, NO |
| 40 | TraesCS2B03G0097700 | 2B | B | Carboxymethylenebutenolidase 1 | LRL | CS, MA, ST, NO, JA |
| 41 | TraesCS6A03G0042300 | 6A | A | RNA binding | MA, LRL | CS, ST, NO, JA |
| 42 | TraesCS2B03G0935600 | 2B | B | Kelch repeat-containing protein | LRL, ST | CS, MA, NO, JA |
| 43 | TraesCS2B03G0919400 | 2B | B | 3-isopropylmalate dehydrogenase | LRL | CS, MA, ST, NO, JA |
| 44 | TraesCS2B03G1103700 | 2B | B | Acetylglutamate kinase-like protein | LRL | CS, MA, ST, NO, JA |
| 45 | TraesCS1B03G0874400 | 1B | B | Proteasome subunit alpha type | MA, LRL, NO | CS, ST, JA |
| 46 | TraesCS2B03G0711300 | 2B | B | E3 ubiquitin-protein ligase family | LRL | CS, MA, ST, NO, JA |
| 47 | TraesCS2B03G1127800 | 2B | B | UDP-N-acetylglucosamine pyrophosphorylas | LRL | CS, MA, ST, NO, JA |
| 48 | TraesCS1B03G0517800 | 1B | B | Coatomer subunit gamma | MA, ST, JA | CS, LRL, NO |
| 49 | TraesCS1B03G0760700 | 1B | B | — | MA | CS, LRL, ST, NO, JA |
| 50 | TraesCS1B03G0729200 | 1B | B | Rhomboid family protein | MA, JA | CS, LRL, ST, NO |
| 51 | TraesCS7B03G0788500 | 7B | B | Receptor-like protein kinase | LRL, ST, NO | CS, MA, JA |
| 52 | TraesCS2B03G1119500 | 2B | B | Aminotransferase | LRL | CS, MA, ST, NO, JA |
| 53 | TraesCS7A03G0259600 | 7A | A | — | JA | CS, MA, LRL, ST, NO |
| 54 | TraesCS5B03G0834800 | 5B | B | WD-repeat protein, putative | ST, JA | CS, MA, LRL, NO |
| 55 | TraesCS2B03G0357400 | 2B | B | — | LRL | CS, MA, ST, NO, JA |
| 56 | TraesCS6B03G0027900 | 6B | B | Guanine nucleotide-binding protein subun | LRL | CS, MA, ST, NO, JA |
| 57 | TraesCS2B03G1408600 | 2B | B | Diacylglycerol kinase | LRL | CS, MA, ST, NO, JA |
| 58 | TraesCS2B03G0753100 | 2B | B | Basic helix-loop-helix (BHLH) Transcript | LRL | CS, MA, ST, NO, JA |
| 59 | TraesCS2B03G1217500 | 2B | B | UDP-glucose 4-epimerase, putative | LRL | CS, MA, ST, NO, JA |
| 60 | TraesCS2B03G0979500 | 2B | B | Proliferating cell nuclear antigen | ST, JA | CS, MA, LRL, NO |
| 61 | TraesCS2B03G1439700 | 2B | B | Calreticulin-like protein | ST, JA | CS, MA, LRL, NO |
| 62 | TraesCS7A03G1301800 | 7A | A | RING/U-box superfamily protein | MA, LRL | CS, ST, NO, JA |
| 63 | TraesCS7B03G1028500 | 7B | B | Copper-transporting ATPase | MA, LRL | CS, ST, NO, JA |
| 64 | TraesCS7B03G0245500 | 7B | B | Sphingoid base hydroxylase 2 | MA, LRL | CS, ST, NO, JA |
| 65 | TraesCS2B03G0894300 | 2B | B | 14-3-3 protein | LRL | CS, MA, ST, NO, JA |
| 66 | TraesCS3B03G0153600 | 3B | B | 14-3-3 protein | NO | CS, MA, LRL, ST, JA |
| 67 | TraesCS7B03G1124800 | 7B | B | WRKY transcription factor | MA, ST, JA | CS, LRL, NO |
| 68 | TraesCS2B03G1082200 | 2B | B | Calcium-dependent protein kinase | LRL | CS, MA, ST, NO, JA |
| 69 | TraesCS2B03G1205900 | 2B | B | Ring finger protein | LRL | CS, MA, ST, NO, JA |
| 70 | TraesCS3B03G0668400 | 3B | B | Cytochrome P450 | MA, LRL, ST, JA | CS, NO |
| 71 | TraesCS6D03G0538500 | 6D | D | MYB transcription factor | MA | CS, LRL, ST, NO, JA |
| 72 | TraesCS2B03G1070700 | 2B | B | Dof zinc finger protein | LRL | CS, MA, ST, NO, JA |
| 73 | TraesCS1D03G0773200 | 1D | D | — | MA, LRL, ST, NO, JA | CS |
| 74 | TraesCS6B03G0478800 | 6B | B | Phosphoglycerate kinase | ST, JA | CS, MA, LRL, NO |

Table S9. Complete list of candidate loci for the dominance prediction (42 groups, 42 candidate homoeologs). Conventions as in Table S7. In none of these does the inserted homoeolog reach the dominant category in every carrier; in 29 of 42 it is balanced in carriers while suppressed in non-carriers, so this class represents the absence of suppression rather than elevated expression. Approximately 40% of this list is expected by chance.

| **No.** | **Gene (v2.1)** | **Chr** | **Sub** | **Annotation** | **TE present in** | **TE absent in** |
| --- | --- | --- | --- | --- | --- | --- |
| 1 | TraesCS2B03G1482800 | 2B | B | Elongation factor 1-alpha | CS, LRL, ST, JA | MA, NO |
| 2 | TraesCS2A03G0005800 | 2A | A | — | CS, LRL, NO | MA, ST, JA |
| 3 | TraesCS7D03G0131300 | 7D | D | SPX domain-containing family protein | NO | CS, MA, LRL, ST, JA |
| 4 | TraesCS7B03G1171400 | 7B | B | XH/XS domain-containing family protein | CS, MA, ST, NO | LRL, JA |
| 5 | TraesCS3D03G1188800 | 3D | D | — | CS, MA | LRL, ST, NO, JA |
| 6 | TraesCS4D03G0029000 | 4D | D | — | CS, LRL, NO, JA | MA, ST |
| 7 | TraesCS3B03G0900700 | 3B | B | Pentatricopeptide repeat-containing prot | CS, LRL, ST | MA, NO, JA |
| 8 | TraesCS4B03G0592100 | 4B | B | — | ST | CS, MA, LRL, NO, JA |
| 9 | TraesCS5A03G0749000 | 5A | A | PHD-finger family protein, expressed | JA | CS, MA, LRL, ST, NO |
| 10 | TraesCS7A03G0855700 | 7A | A | — | CS, NO | MA, LRL, ST, JA |
| 11 | TraesCS2D03G0479800 | 2D | D | — | LRL | CS, MA, ST, NO, JA |
| 12 | TraesCS1B03G0966900 | 1B | B | Peptidyl-tRNA hydrolase, putative | CS, LRL, NO, JA | MA, ST |
| 13 | TraesCS1A03G0958900 | 1A | A | 3-deoxy-manno-octulosonate cytidylyltran | CS, LRL, ST, NO, JA | MA |
| 14 | TraesCS2B03G0727100 | 2B | B | Aminotransferase | CS, MA, ST, NO, JA | LRL |
| 15 | TraesCS6D03G0889800 | 6D | D | — | JA | CS, MA, LRL, ST, NO |
| 16 | TraesCS5D03G1219200 | 5D | D | — | CS, NO | MA, LRL, ST, JA |
| 17 | TraesCS1D03G1003400 | 1D | D | Protein N-methyltransferase | CS, MA, LRL, ST, NO | JA |
| 18 | TraesCS2D03G0238100 | 2D | D | — | LRL | CS, MA, ST, NO, JA |
| 19 | TraesCS2B03G0904800 | 2B | B | ATP-dependent 6-phosphofructokinase | CS, MA, ST, NO, JA | LRL |
| 20 | TraesCS2A03G0047100 | 2A | A | ATP-dependent Clp protease adapter prote | CS, LRL, NO | MA, ST, JA |
| 21 | TraesCS5B03G0754000 | 5B | B | — | CS, MA, LRL, ST, JA | NO |
| 22 | TraesCS2B03G0725100 | 2B | B | Aspartyl/glutamyl-tRNA(Asn/Gln) amidotra | CS, MA, ST, NO, JA | LRL |
| 23 | TraesCS2B03G0600200 | 2B | B | Trafficking protein particle complex sub | CS, MA, ST, NO, JA | LRL |
| 24 | TraesCS2B03G0919500 | 2B | B | — | CS, MA, ST, NO, JA | LRL |
| 25 | TraesCS2B03G0845600 | 2B | B | — | CS, MA, ST, NO, JA | LRL |
| 26 | TraesCS2B03G0811400 | 2B | B | Cell division protein ftsZ | CS, MA, ST, NO, JA | LRL |
| 27 | TraesCS2B03G1055300 | 2B | B | Mechanosensitive ion channel protein | CS, MA, ST, NO, JA | LRL |
| 28 | TraesCS4B03G0004300 | 4B | B | Phosphate transporter | CS, MA, ST, NO, JA | LRL |
| 29 | TraesCS5B03G1028400 | 5B | B | Calmodulin-binding protein, putative, ex | CS, ST, NO | MA, LRL, JA |
| 30 | TraesCS2B03G0388800 | 2B | B | Transmembrane protein 115 | CS, MA, ST, NO, JA | LRL |
| 31 | TraesCS5A03G0143000 | 5A | A | Cytochrome P450, putative | CS, NO, JA | MA, LRL, ST |
| 32 | TraesCS5B03G0715800 | 5B | B | N-methyl-L-tryptophan oxidase | ST | CS, MA, LRL, NO, JA |
| 33 | TraesCS1D03G1004900 | 1D | D | Acyl-protein thioesterase 2 | CS, MA, LRL, ST, NO | JA |
| 34 | TraesCS3B03G1380100 | 3B | B | — | CS, MA, LRL, NO, JA | ST |
| 35 | TraesCS4B03G0039300 | 4B | B | Ras family protein | CS, LRL, NO, JA | MA, ST |
| 36 | TraesCS2B03G0740100 | 2B | B | Glutamate receptor | CS, MA, ST, NO, JA | LRL |
| 37 | TraesCS2B03G0271900 | 2B | B | transcription factor-like protein | CS, LRL, ST, NO, JA | MA |
| 38 | TraesCS2B03G0779300 | 2B | B | Acyl-[acyl-carrier-protein] desaturase | CS, MA, ST, NO, JA | LRL |
| 39 | TraesCS1B03G0668700 | 1B | B | Sugar transporter, putative | MA, ST, JA | CS, LRL, NO |
| 40 | TraesCS1D03G1048400 | 1D | D | Histone H1 | CS, MA, LRL, ST, NO | JA |
| 41 | TraesCS7B03G0545300 | 7B | B | Chloroplast envelope membrane protein | CS, MA, ST, NO, JA | LRL |
| 42 | TraesCS4B03G0398400 | 4B | B | — | CS, LRL, ST, NO, JA | MA |

Table S10. Candidate loci from the pan-transcriptome of White et al. (2025), both directions (77 suppression and 106 dominance candidate homoeologs). Tissue codes: D, aerial organs at dawn; V, aerial organs at dusk; R, root; S, spike; F, flag leaf; G, grain. Cultivars: ARI, ArinaLrFor; JAG, Jagger; JUL, Julius; LAC, LongReach Lancer; LAN, CDC Landmark; MAC, Mace; NOR, Norin 61; STA, CDC Stanley; SYM, SY Mattis. These derive from the authors' published bias classes, which were called against Chinese Spring.

| **No.** | **Direction** | **CS orthologue (v2.1)** | **Sub** | **Tissue(s)** | **Annotation** | **TE present in** |
| --- | --- | --- | --- | --- | --- | --- |
| 1 | Suppression | TraesCS3B03G0614600 | B | DVRSFG | Mitochondrial carrier protein-like | ARI, JAG, JUL, LAC, LAN, MAC, STA, SYM |
| 2 | Suppression | — | D | DVRSF | — | ARI, JAG, JUL, LAC, LAN, NOR, STA, SYM |
| 3 | Suppression | TraesCS2A03G0487500 | A | RSFG | Ribosomal rna small subunit methyltr | ARI, JUL, LAC, LAN, MAC, STA, SYM |
| 4 | Suppression | TraesCS5D03G0919800 | D | DVRS | Alpha/beta-Hydrolases superfamily pr | JAG, SYM |
| 5 | Suppression | TraesCS2B03G0525600 | B | RSG | — | LAC |
| 6 | Suppression | TraesCS3A03G0639900 | A | DVF | Peroxidase | STA |
| 7 | Suppression | TraesCS6A03G0048200 | A | RSG | Mitochondrial inner membrane proteas | ARI, LAC, MAC |
| 8 | Suppression | TraesCS1B03G0107600 | B | DV | 3-ketoacyl-CoA synthase | NOR |
| 9 | Suppression | TraesCS1D03G0546300 | D | DF | Kinase like protein | JAG, MAC |
| 10 | Suppression | TraesCS1D03G0673100 | D | DG | Adenine nucleotide alpha hydrolases- | ARI, JAG, JUL, STA, SYM |
| 11 | Suppression | TraesCS2A03G0179800 | A | RF | — | SYM |
| 12 | Suppression | TraesCS2D03G0240500 | D | SG | — | JAG |
| 13 | Suppression | TraesCS2B03G0467200 | B | DF | Aminotransferase | LAN |
| 14 | Suppression | TraesCS2B03G0719400 | B | DS | L-ascorbate oxidase-like protein | ARI, JUL, LAN, MAC, NOR, STA, SYM |
| 15 | Suppression | TraesCS2D03G0598100 | D | DS | L-ascorbate oxidase-like protein | ARI, JAG, LAC, LAN, MAC, NOR, STA, SYM |
| 16 | Suppression | TraesCS2B03G1043500 | B | DS | Cyclin-like | LAC |
| 17 | Suppression | TraesCS2B03G1045200 | B | DV | Blue copper protein | LAN, MAC |
| 18 | Suppression | TraesCS2D03G1091300 | D | DF | Cell division protein ftsZ | ARI, JAG, JUL, SYM |
| 19 | Suppression | — | A | DV | — | MAC |
| 20 | Suppression | TraesCS4D03G0179600 | D | DF | — | ARI, JUL, NOR |
| 21 | Suppression | TraesCS5B03G0234000 | B | DV | Pectinesterase | SYM |
| 22 | Suppression | TraesCS5B03G0447000 | B | VR | — | JUL |
| 23 | Suppression | — | B | DR | — | SYM |
| 24 | Suppression | — | D | V | — | ARI, JAG, JUL, LAC, LAN, MAC, NOR, SYM |
| 25 | Suppression | TraesCS1A03G0823700 | A | R | — | NOR |
| 26 | Suppression | TraesCS1B03G1195400 | B | R | — | MAC |
| 27 | Suppression | TraesCS2A03G0021300 | A | R | — | ARI, JAG, JUL, LAN, MAC, NOR, STA, SYM |
| 28 | Suppression | TraesCS2B03G0312500 | B | R | Peroxiredoxin | LAC |
| 29 | Suppression | TraesCS2B03G0450300 | B | F | Heavy-metal-associated domain-contai | JUL |
| 30 | Suppression | TraesCS2A03G0402800 | A | G | Pentatricopeptide repeat-containing | JUL |
| 31 | Suppression | TraesCS2B03G0514400 | B | G | Pentatricopeptide repeat-containing | JUL |
| 32 | Suppression | TraesCS2D03G0404600 | D | G | Pentatricopeptide repeat-containing | JUL, SYM |
| 33 | Suppression | TraesCS2B03G0568900 | B | F | Serine/threonine-protein kinase | NOR |
| 34 | Suppression | TraesCS2B03G0591600 | B | S | WAT1-related protein | LAC |
| 35 | Suppression | — | B | R | — | LAC |
| 36 | Suppression | TraesCS2B03G0856800 | B | G | MLO-like protein | LAC |
| 37 | Suppression | TraesCS2B03G0880400 | B | D | Sugar transporter, putative | LAC |
| 38 | Suppression | TraesCS2B03G1109100 | B | S | Nudix hydrolase | LAC |
| 39 | Suppression | TraesCS2D03G1064800 | D | R | — | LAC, LAN, MAC, NOR, STA |
| 40 | Suppression | — | B | R | — | LAC |
| 41 | Suppression | TraesCS2A03G1246800 | A | G | Gibberellin 3-beta-hydroxylase 2 | LAC |
| 42 | Suppression | TraesCS3A03G0040500 | A | S | WAT1-related protein | LAN |
| 43 | Suppression | TraesCS3B03G0060700 | B | S | WAT1-related protein | LAN |
| 44 | Suppression | TraesCS3D03G0048300 | D | S | WAT1-related protein | ARI, JAG, JUL, LAC, LAN, MAC, NOR, STA |
| 45 | Suppression | TraesCS3A03G0358200 | A | S | — | JUL |
| 46 | Suppression | TraesCS3B03G0668400 | B | F | Cytochrome P450 | JAG, JUL, LAC, LAN, MAC, STA, SYM |
| 47 | Suppression | TraesCS3B03G0756000 | B | F | Receptor-like protein kinase | JAG, JUL, LAC, LAN, MAC, NOR, STA, SYM |
| 48 | Suppression | TraesCS3D03G0605000 | D | F | Receptor-like protein kinase | ARI |
| 49 | Suppression | TraesCS3A03G0664000 | A | F | — | ARI, JUL, LAC, LAN, NOR, STA, SYM |
| 50 | Suppression | — | B | G | — | SYM |
| 51 | Suppression | TraesCS3B03G0811300 | B | R | GDSL esterase/lipase | ARI, JUL, LAC, LAN, MAC, NOR, STA, SYM |
| 52 | Suppression | TraesCS3B03G0815700 | B | R | — | SYM |
| 53 | Suppression | TraesCS3B03G0876900 | B | G | Cytokinin oxidase/dehydrogenase | SYM |
| 54 | Suppression | — | D | F | — | ARI, JAG, LAN, NOR, STA |
| 55 | Suppression | — | D | S | — | LAC |
| 56 | Suppression | TraesCS4B03G0557600 | B | F | Glutathione-S-transferase | STA |
| 57 | Suppression | TraesCS4B03G0172600 | B | D | — | JAG, JUL, STA, SYM |
| 58 | Suppression | TraesCS4B03G0146400 | B | G | Dimeric alpha-amylase inhibitor | NOR, SYM |
| 59 | Suppression | — | B | G | — | ARI, JAG, LAC, LAN, NOR, SYM |
| 60 | Suppression | — | D | G | — | ARI, JAG, JUL, LAC, NOR |
| 61 | Suppression | TraesCS5B03G1263200 | B | G | Zinc finger family protein | NOR |
| 62 | Suppression | — | D | S | — | NOR |
| 63 | Suppression | TraesCS5D03G0417500 | D | R | — | NOR |
| 64 | Suppression | TraesCS5A03G0566000 | A | V | Tetraspanin family protein | JAG |
| 65 | Suppression | TraesCS5A03G0661500 | A | V | Cinnamoyl-CoA reductase 4 | ARI, JAG, LAC, LAN, MAC, STA |
| 66 | Suppression | TraesCS5A03G0723800 | A | R | ENTH/ANTH/VHS superfamily protein | NOR |
| 67 | Suppression | TraesCS5B03G0954100 | B | V | IQ domain-containing protein | NOR |
| 68 | Suppression | TraesCS5A03G0968300 | A | R | Leucine carboxyl methyltransferase | ARI, JUL, LAC, LAN, MAC, NOR, STA, SYM |
| 69 | Suppression | TraesCS6B03G0318400 | B | V | Kinase-like protein | MAC |
| 70 | Suppression | TraesCS6D03G0288600 | D | S | Pentatricopeptide repeat-containing | NOR |
| 71 | Suppression | TraesCS6B03G0464700 | B | R | Serine-threonine protein phosphatase | STA |
| 72 | Suppression | TraesCS6A03G0831000 | A | F | — | STA |
| 73 | Suppression | TraesCS7A03G0485700 | A | V | — | NOR |
| 74 | Suppression | TraesCS7B03G0425300 | B | G | F-box protein | ARI, JAG, JUL, LAN, NOR, SYM |
| 75 | Suppression | TraesCS7A03G0764600 | A | R | Zinc transport protein ZntB | SYM |
| 76 | Suppression | TraesCS7A03G0786700 | A | G | — | SYM |
| 77 | Suppression | TraesCS7B03G0682200 | B | G | Glycosyltransferase | NOR |
| 1 | Dominance | TraesCS1B03G0254600 | B | G | — | JUL |
| 2 | Dominance | TraesCS1A03G0193800 | A | V | — | JAG, JUL, LAC, MAC, NOR, STA, SYM |
| 3 | Dominance | TraesCS1A03G0308600 | A | G | Peroxidase | ARI, JAG, JUL, LAC, MAC, STA, SYM |
| 4 | Dominance | TraesCS1B03G0390800 | B | G | Peroxidase | ARI, JAG, JUL, LAC, MAC, NOR, STA |
| 5 | Dominance | TraesCS1D03G0310800 | D | G | Peroxidase | LAC |
| 6 | Dominance | TraesCS1B03G0482100 | B | D | — | JUL |
| 7 | Dominance | TraesCS1D03G0364900 | D | D | — | ARI, JAG, JUL, LAC, MAC, NOR, STA |
| 8 | Dominance | TraesCS1B03G0672600 | B | S | — | NOR |
| 9 | Dominance | — | A | F | — | JUL, LAC, STA |
| 10 | Dominance | TraesCS1D03G0728500 | D | D | — | NOR |
| 11 | Dominance | — | B | R | — | ARI |
| 12 | Dominance | TraesCS1B03G1195600 | B | G | Auxin efflux carrier component | JUL, LAC, MAC, NOR, STA, SYM |
| 13 | Dominance | TraesCS2A03G0066300 | A | G | Nicotianamine synthase | JUL |
| 14 | Dominance | TraesCS2A03G0084800 | A | G | — | ARI, JUL, LAC, MAC, NOR, STA, SYM |
| 15 | Dominance | TraesCS2A03G0229600 | A | V | Kelch repeat-containing protein | ARI, JAG, JUL, LAC, MAC, STA, SYM |
| 16 | Dominance | TraesCS2B03G0504800 | B | DVG | Receptor kinase | LAC |
| 17 | Dominance | TraesCS2B03G0533100 | B | DVG | Glucan endo-1,3-beta-glucosidase, pu | ARI, JAG, JUL, MAC, NOR, STA, SYM |
| 18 | Dominance | TraesCS2A03G0418400 | A | S | — | JAG |
| 19 | Dominance | TraesCS2B03G0590600 | B | V | — | LAC |
| 20 | Dominance | TraesCS2B03G0695400 | B | DVRSF | — | JAG, JUL, MAC, NOR, STA |
| 21 | Dominance | TraesCS2B03G0781000 | B | G | Small ribosomal subunit biogenesis G | ARI, JAG, JUL, MAC, NOR, STA, SYM |
| 22 | Dominance | TraesCS2B03G0858800 | B | F | Ferric reduction oxidase 7 | ARI, JAG, JUL, MAC, NOR, STA, SYM |
| 23 | Dominance | — | A | DVRF | — | NOR |
| 24 | Dominance | TraesCS2B03G0860500 | B | G | — | ARI, JAG, JUL, MAC, NOR, STA, SYM |
| 25 | Dominance | TraesCS2D03G0761700 | D | S | WRKY transcription factor | ARI, JUL, LAC, MAC, NOR, STA, SYM |
| 26 | Dominance | TraesCS2B03G0882300 | B | DV | Sugar transporter, putative | ARI, JAG, JUL, MAC, NOR, STA, SYM |
| 27 | Dominance | TraesCS2A03G0891700 | A | SG | — | ARI, JAG, LAC, MAC, NOR, STA, SYM |
| 28 | Dominance | TraesCS2B03G0984800 | B | SG | — | ARI, JAG, JUL, MAC, NOR, STA, SYM |
| 29 | Dominance | TraesCS2A03G0897800 | A | V | — | JAG, JUL, LAC, MAC, NOR |
| 30 | Dominance | TraesCS2B03G0988500 | B | V | — | ARI, JAG, JUL, MAC, NOR, STA, SYM |
| 31 | Dominance | TraesCS2B03G1026200 | B | D | Acid invertase 1 | LAC |
| 32 | Dominance | TraesCS2A03G0941200 | A | G | Histone H1 | LAC, MAC |
| 33 | Dominance | TraesCS2B03G1041800 | B | D | Subtilisin-like protease | LAC |
| 34 | Dominance | TraesCS2B03G1217100 | B | S | — | JUL, LAC |
| 35 | Dominance | TraesCS2B03G1220200 | B | V | — | MAC |
| 36 | Dominance | TraesCS3B03G0103500 | B | G | Calmodulin-binding protein-like | JUL |
| 37 | Dominance | TraesCS3A03G0188600 | A | V | Pentatricopeptide repeat-containing | ARI, JAG, JUL, LAC, MAC, STA, SYM |
| 38 | Dominance | TraesCS3D03G0167800 | D | V | Pentatricopeptide repeat-containing | ARI, JUL, STA |
| 39 | Dominance | — | B | DVSF | — | MAC |
| 40 | Dominance | TraesCS3B03G0398000 | B | D | Cationic amino acid transporter | JUL |
| 41 | Dominance | TraesCS3D03G0316300 | D | D | Cationic amino acid transporter | ARI, JAG, JUL, MAC, STA, SYM |
| 42 | Dominance | TraesCS3B03G0403500 | B | D | — | ARI, JUL, STA, SYM |
| 43 | Dominance | TraesCS3D03G0319100 | D | D | — | ARI, JAG, LAC, MAC, NOR |
| 44 | Dominance | TraesCS3B03G0434500 | B | V | Auxin-responsive protein | JAG, JUL, LAC, MAC, NOR, STA, SYM |
| 45 | Dominance | TraesCS3B03G0514300 | B | DVRS | Translin family protein | JAG, JUL, LAC, MAC, NOR, STA, SYM |
| 46 | Dominance | TraesCS3D03G0504100 | D | DVRSF | Protein phosphatase 2C | ARI, JUL, LAC, MAC, NOR, STA, SYM |
| 47 | Dominance | TraesCS3A03G0565500 | A | S | Pentatricopeptide repeat-containing | ARI, JUL, LAC, MAC, SYM |
| 48 | Dominance | — | B | G | — | NOR |
| 49 | Dominance | — | D | G | — | STA |
| 50 | Dominance | — | B | R | — | ARI, JAG, JUL, LAC, NOR, STA, SYM |
| 51 | Dominance | TraesCS3B03G0707900 | B | DRS | Homeobox protein, putative | ARI, JAG, JUL, LAC, NOR, STA, SYM |
| 52 | Dominance | TraesCS3A03G0664000 | A | G | — | ARI, JUL, LAC, NOR, STA, SYM |
| 53 | Dominance | TraesCS3B03G0841500 | B | G | — | ARI, JAG, JUL, LAC, NOR |
| 54 | Dominance | TraesCS3B03G0912100 | B | F | Purple acid phosphatase | ARI, JUL, NOR, SYM |
| 55 | Dominance | TraesCS3B03G0928200 | B | DVRSFG | — | JAG, LAC, MAC, STA, SYM |
| 56 | Dominance | TraesCS3B03G0948900 | B | V | — | STA |
| 57 | Dominance | TraesCS3D03G0915600 | D | R | Selenium-binding protein | JUL, MAC, NOR, SYM |
| 58 | Dominance | TraesCS4A03G0042200 | A | S | Annexin | MAC |
| 59 | Dominance | TraesCS4A03G0083100 | A | DVRSFG | — | LAC, MAC, NOR |
| 60 | Dominance | TraesCS4B03G0656900 | B | R | Receptor-like kinase | ARI, JAG, JUL, MAC, NOR, STA, SYM |
| 61 | Dominance | TraesCS4B03G0630500 | B | DVR | — | ARI, LAC, MAC, STA, SYM |
| 62 | Dominance | TraesCS4A03G0278400 | A | R | NAC domain protein | JAG, NOR |
| 63 | Dominance | TraesCS4A03G0481200 | A | R | Glucan endo-1,3-beta-glucosidase | JAG |
| 64 | Dominance | — | B | DV | — | JUL, MAC, SYM |
| 65 | Dominance | TraesCS4A03G0849800 | A | F | Pentatricopeptide repeat-containing | ARI, JAG, SYM |
| 66 | Dominance | TraesCS4D03G0720800 | D | S | — | ARI, JUL, LAC, MAC, NOR, STA, SYM |
| 67 | Dominance | — | A | VR | — | ARI, JAG, JUL, MAC, NOR, STA, SYM |
| 68 | Dominance | — | B | VS | — | MAC |
| 69 | Dominance | — | B | D | — | JUL, STA |
| 70 | Dominance | TraesCS5B03G0412100 | B | R | — | ARI, JAG, JUL, LAC, STA, SYM |
| 71 | Dominance | TraesCS5B03G0493700 | B | D | — | ARI, JAG, LAC, NOR, STA, SYM |
| 72 | Dominance | TraesCS5A03G0513200 | A | DVSG | Pentatricopeptide repeat-containing | MAC |
| 73 | Dominance | TraesCS5D03G0479300 | D | DVSG | Heat shock factor-binding protein 1 | JUL, STA |
| 74 | Dominance | TraesCS5A03G0603600 | A | DRSFG | Calcium-binding protein | ARI, JAG, JUL, LAC, NOR, STA |
| 75 | Dominance | TraesCS5B03G0622300 | B | DRSFG | Calcium-binding protein | JAG, JUL, MAC, NOR, STA, SYM |
| 76 | Dominance | TraesCS5D03G0569100 | D | DRSFG | Calcium-binding protein | MAC, NOR, STA, SYM |
| 77 | Dominance | TraesCS5A03G0610500 | A | D | Alpha/beta-Hydrolases superfamily pr | NOR |
| 78 | Dominance | — | A | D | — | JUL |
| 79 | Dominance | — | B | D | — | SYM |
| 80 | Dominance | TraesCS5D03G0878300 | D | DSF | General transcription factor 3C poly | ARI, JUL, LAC, MAC, NOR, STA, SYM |
| 81 | Dominance | TraesCS5B03G1011700 | B | G | RNA-binding protein-like | NOR |
| 82 | Dominance | TraesCS5B03G1047400 | B | D | FAD/NAD(P)-binding oxidoreductase fa | SYM |
| 83 | Dominance | TraesCS5D03G1065200 | D | DR | C2 calcium/lipid-binding and GRAM do | ARI, JAG, LAC, MAC, NOR, STA, SYM |
| 84 | Dominance | TraesCS5B03G1202900 | B | V | Ribosomal RNA large subunit methyltr | LAC |
| 85 | Dominance | TraesCS5D03G1090300 | D | V | Urease subunit alpha | ARI, JAG, JUL, LAC, NOR, STA, SYM |
| 86 | Dominance | TraesCS4D03G0761300 | D | DVRSF | F-box/LRR-repeat protein | ARI, JAG, JUL, LAC, NOR, STA, SYM |
| 87 | Dominance | TraesCS6A03G0401800 | A | V | Glucan endo-1,3-beta-glucosidase | STA |
| 88 | Dominance | TraesCS6D03G0351300 | D | DV | — | LAC |
| 89 | Dominance | — | D | S | — | MAC |
| 90 | Dominance | TraesCS6B03G0889800 | B | S | Rho GDP-dissociation inhibitor 1 | ARI, JAG, JUL, MAC, NOR, STA, SYM |
| 91 | Dominance | TraesCS7B03G0200400 | B | VRSG | WAT1-related protein | NOR |
| 92 | Dominance | — | D | DRS | — | JUL, LAC |
| 93 | Dominance | TraesCS7D03G0494800 | D | R | — | JAG |
| 94 | Dominance | TraesCS7A03G0635400 | A | D | MYB transcription factor | NOR |
| 95 | Dominance | — | B | D | — | MAC, NOR |
| 96 | Dominance | TraesCS7B03G0575000 | B | F | Pentatricopeptide repeat-containing | ARI, JAG, MAC, STA, SYM |
| 97 | Dominance | TraesCS7D03G0765200 | D | S | Ferredoxin | JUL |
| 98 | Dominance | TraesCS7A03G0909500 | A | FG | Protein OVEREXPRESSOR OF CATIONIC PE | JUL |
| 99 | Dominance | TraesCS7D03G0941700 | D | R | Pentatricopeptide repeat-containing | ARI, LAC, SYM |
| 100 | Dominance | TraesCS7A03G0982800 | A | S | Eukaryotic aspartyl protease family | ARI, JAG, JUL, LAC, MAC, NOR, SYM |
| 101 | Dominance | TraesCS7B03G0960400 | B | D | Serine/threonine-protein kinase | ARI, JAG, LAC, MAC, NOR, STA, SYM |
| 102 | Dominance | — | D | SFG | — | ARI, JUL, LAC, MAC, NOR, STA, SYM |
| 103 | Dominance | TraesCS7B03G1267400 | B | G | ATP-dependent zinc metalloprotease F | ARI, JAG, JUL, LAC, STA, SYM |
| 104 | Dominance | TraesCS7B03G1274100 | B | S | — | JUL |
| 105 | Dominance | TraesCS7D03G1265000 | D | S | — | ARI |
| 106 | Dominance | TraesCS7B03G1304400 | B | DVRSFG | — | ARI, JAG, JUL, LAC, MAC, STA, SYM |
